# Structural surfaceomics reveals an AML-specific conformation of Integrin-β2 as a CAR-T therapy target

**DOI:** 10.1101/2022.10.10.511511

**Authors:** Kamal Mandal, Gianina Wicaksono, Clinton Yu, Jarrett J. Adams, Michael R. Hoopmann, William C. Temple, Bonell Patiño Escobar, Maryna Gorelik, Christian H. Ihling, Matthew A. Nix, Akul Naik, Emilio Ramos, Corynn Kasap, Veronica Steri, Juan Antonio Camara Serrano, Fernando Salangsang, Paul Phojanakong, Melanie McMillan, Victor Gavallos, Andrew D. Leavitt, Andrea Sinz, Benjamin J. Huang, Elliot Stieglitz, Catherine C. Smith, Robert L. Moritz, Sachdeva S. Sidhu, Lan Huang, Arun P. Wiita

**Affiliations:** Department of Laboratory Medicine, University of California, San Francisco, CA, USA; Dept. of Pediatrics, Division of Hematology/Oncology, University of California, San Francisco, CA, USA; Department of Physiology & Biophysics, University of California, Irvine, CA, USA; The Donnelly Centre, University of Toronto, ON, Canada; Dept. of Medicine, Division of Hematology/Oncology, University of California, San Francisco, CA, USA; Department of Pharmaceutical Chemistry & Bioanalytics, Institute of Pharmacy, Martin-Luther University Halle-Wittenberg, Halle, Germany; Institute for Systems Biology, Seattle, WA, USA; Helen Diller Family Comprehensive Cancer Center, University of California, San Francisco, CA, USA; Department of Pediatrics, Division of Allergy, Immunology, and Bone Marrow Transplantation, University of California, San Francisco, San Francisco, CA

**Keywords:** Proteomics, XL-MS, acute myeloid leukemia, CAR-T, immunotherapy

## Abstract

Safely expanding indications for cellular therapies has been challenging given a lack of highly cancer-specific surface markers. Here, we explore the hypothesis that tumor cells express cancer-specific surface protein conformations, invisible to standard target discovery pipelines evaluating gene or protein expression, that can be identified and immunotherapeutically targeted. We term this strategy, integrating cross-linking mass spectrometry (XL-MS) with glycoprotein surface capture, “structural surfaceomics”. As a proof of principle, we apply this technology to acute myeloid leukemia, a hematologic malignancy with dismal outcomes and no known optimal immunotherapy target. We identify the activated conformation of integrin-β2 as a structurally-defined, widely-expressed, AML-specific target. We develop and characterize recombinant antibodies to this protein conformation, and show that chimeric antigen receptor (CAR) T-cells eliminate AML cells and patient-derived xenografts without notable toxicity versus normal hematopoietic cells. Our findings validate an AML conformation-specific target antigen while demonstrating a toolkit for applying these strategies more broadly.

## INTRODUCTION

Cellular therapies are one of the most exciting modalities in cancer care, leading to the promise of long-term tumor control as “living drugs”^1^. However, safely applying these therapies to cancers beyond B-cell malignancies has remained clinically challenging^2^. A major hurdle remains identification of surface antigens that are specifically expressed on tumor cells but not on other essential tissues, with a goal of minimizing “on target, off tumor” toxicity^3, 4^.

Recently, we were intrigued by the discovery of an activated conformation of integrin-β7 as a specific cellular therapy in multiple myeloma^5^. In commonly-used target discovery pipelines, relying entirely on analysis of transcript and/or protein expression levels^6^, integrin-β7 would not be considered an optimal target due to widespread expression on other hematopoietic cells^7^. However, oncogenic signaling was proposed to drive the aberrant constitutive activation of this integrin on myeloma^8, 9^. This change in protein state led to the opportunity to target the active conformation of integrin-β7 while sparing other normal blood cells, where it remained in the closed, resting conformation.

This finding raised the exciting hypothesis that given aberrancies in tumor signaling, metabolism, or cell-microenvironment communication – all of which heavily involve membrane proteins – cancer-specific surface protein conformations may in fact be widespread. However, this result in myeloma was the serendipitous outcome of a hybridoma screen, without any intention to identify a conformation-selective immunotherapy target. Thus, here we aimed to develop a technology to systematically probe this possible untapped source of tumor-specific surface antigens. Specifically, we took advantage of cross-linking mass spectrometry, commonly known as XL-MS^10^. This technology most commonly employs bifunctional lysine-reactive reagents to define inter- or intra-protein interactions based on identified peptide-peptide cross-links. While XL-MS is most often employed to define protein-protein interactions or structural contraints^10^, this approach can also yield low-resolution structural information for hundreds or thousands of proteins in a sample^11, 12^.

However, one of the major hurdles in XL-MS is the low fraction of cross-linked peptides compared to total peptides in any given sample analyzed by mass spectrometry (MS)^13^. Therefore, to specifically focus on cell surface antigens, we combined XL-MS with cell surface capture (CSC), a method to specifically enrich cell surface N-linked glycoproteins^14^. We and others have used CSC to successfully identify immunotherapy targets based on surface protein abundance^15, 16^. Here, by combining XL-MS and CSC in “structural surfaceomics”, we aim to move to the next level of protein-centric target discovery.

As an initial proof of principle, we apply structural surfaceomics to target discovery in acute myeloid leukemia (AML), a frequently-diagnosed hematologic malignancy with dismal prognosis^17^. In contrast to B-cell acute lymphoblastic leukemia, chimeric antigen receptor (CAR) T cells in AML have generally led to either significant toxicities or disappointing clinical efficacy^18, 19^. As demonstrated in an integrated study of the AML transcriptome and surface proteome^16^, one major hurdle to CAR-T therapy for AML is lack of optimal immunotherapy targets. Leading current targets include CD33 and CD123, both of which are expressed widely on AML blasts but also on normal myeloid cells as well as hematopoietic stem and progenitor cells (HSPCs)^18, 20, 21^. Treatment with these CAR-Ts therefore lead to myeloablation and must be followed by allogeneic stem cell transplantation^18^. Other non-myeloablative targets exist, including CLL-1/CLEC12A, but this antigen is also expressed widely on normal myeloid cells, and thus can still spur toxicities, and also shows significant heterogeneity on patient blasts, potentially leading to reduced efficacy^16^. Thus, there remains a significant need to identify AML-specific cellular therapy targets which may eliminate tumor while sparing normal myeloid cells.

Here, we apply structural surfaceomics to an AML model and identify the activated conformation of integrin-β2 as a promising immunotherapeutic target, expressed widely across cell lines and patient tumors. We develop and characterize humanized recombinant antibodies specific for this activated conformation of this protein. We further demonstrate that CAR-T cells incorporating these recombinant binders are efficacious versus AML models, and, importantly, do not show any evidence of toxicity versus normal hematopoietic cells in a humanized immune system murine model, unlike anti-CD33 CAR-T. Our results validate active integrin-β2 as a promising cellular therapy target in AML with a favorable toxicity profile. In addition, our findings suggest structural surfaceomics as a strategy to unlock a previously unexplored class of immunotherapy targets, invisible to standard discovery strategies.

## RESULTS

### Development and application of the structural surfaceomics technology

Our overall strategy for structural surfaceomics is to first use a bifunctional chemical cross-linker applied to live cells, followed by glycoprotein oxidation and biotinylation using the CSC strategy (**Fig. 1a**). The goal of this strategy is to “freeze” the native protein conformation *in situ*, thereby preserving relevant structural information, followed by streptavidin-based enrichment of surface proteins, to increase MS coverage of our most relevant peptides versus much more abundant intracellular protein cross-links.

**Figure 1.**
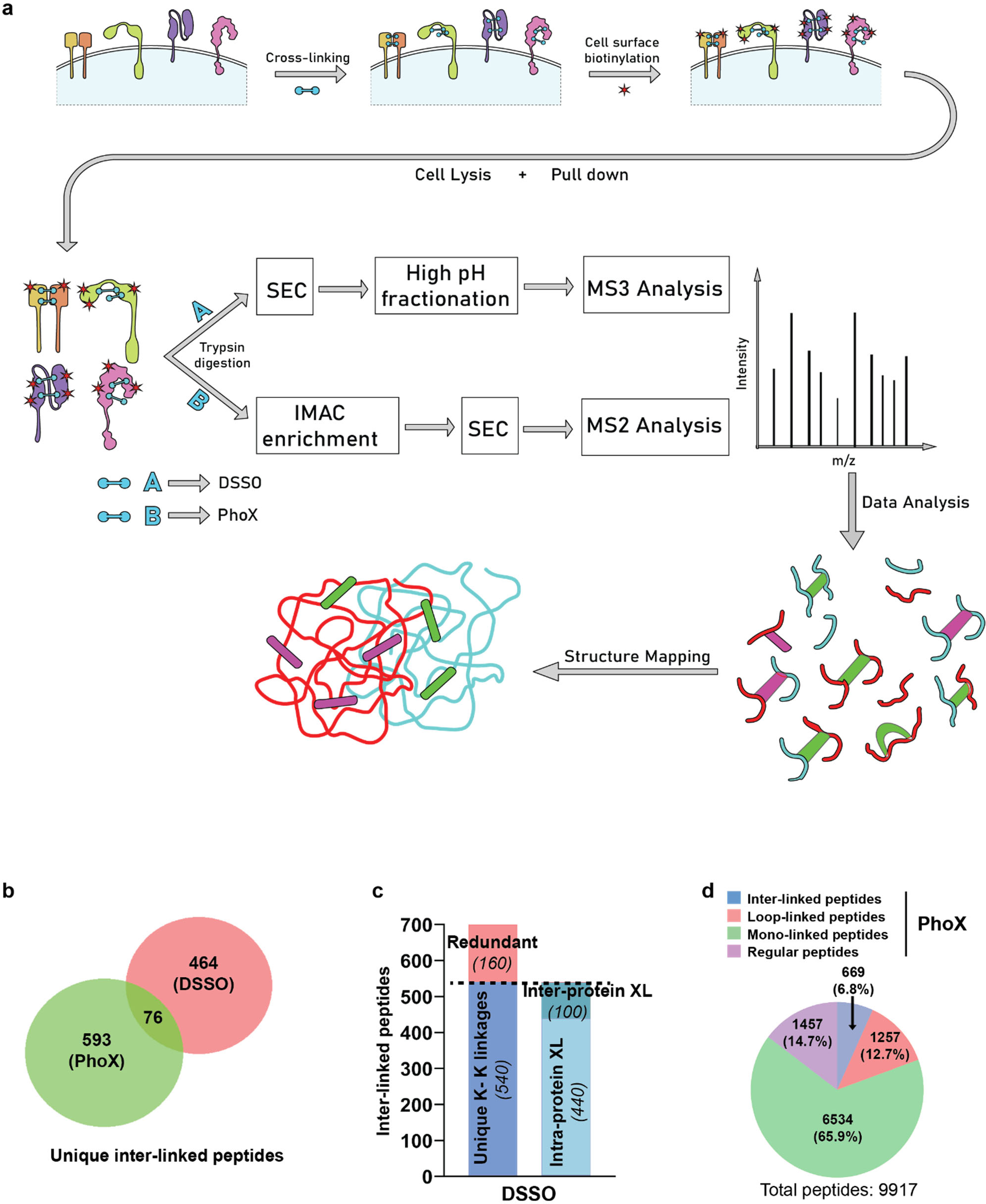
XL-MS + surface glycoprotein capture strategy to identify conformation specific cancer antigens. **a.** Schematic flow diagram of “structural surfaceomics” approach. **b.** Venn diagram showing the total number of cross-linked peptides identified from the two different approaches (MS^2^ and MS^3^ based). PhoX and DSSO was used as a cross-linker for the MS^2^ and MS^3^ approach, respectively. **c.** Bar graph showing distribution of inter- and intra-protein cross-links (XL) from MS^3^ (DSSO) based XL-MS. **d.** Pie chart showing distribution of the various types of cross-links obtained from PhoX MS^2^-based XL-MS. All the cross-links were identified with ≤ 1% FDR (See Methods for details). “Regular” peptides = no PhoX modification detected on any lysines. Source data in Supplementary Dataset 1, 2.

As an initial model system, we used the Nomo-1 AML cell line, derived from a patient with a monocytic leukemia^22^. Using Nomo-1, we explored two complementary chemical strategies in parallel for XL-MS. One strategy incorporates the MS-cleavable cross-linker DSSO (disuccinimidyl sulfoxide), which we and others have used frequently to study protein-protein interactions in both recombinant proteins and whole cell lysates^23–25^. We also employed the recently-described non-cleavable cross-linker PhoX (disuccinimidyl phenyl phosphonic acid) which incorporates a phosphonate-based handle allowing for enrichment of cross-links via immobilized metal affinity chromatography (IMAC)^13^. We applied these strategies in separate experiments to Nomo-1, using cellular input of 0.4-5e9 cells (**Fig. 1a,b**).

XL-MS can identify inter-linked (type 2; bridging two separate peptides), intra-linked (“loop linked”, type 1; two lysines crosslinked in the same peptide), and mono-linked (“dead end”, type 0; single modified lysine) peptides. Inter- and intra-linked peptides could be informative for our strategy, whereas mono-linked are not. For DSSO, we used our previously published XL-MS computational approach^23^ to analyze these data, and also adapted this strategy to a publicly-available version compatible with the Trans-Proteomic Pipeline^26^, called Ving (**Extended Data Fig. 1a, 2** and **Methods**). In our initial DSSO experiment, we enriched crosslinked peptides by size exclusion chromatography (SEC) alone, whereas in our subsequent experiment we followed SEC with tip-based reversed-phase high pH fractionation (HpHt) to optimize coverage^27^. Between these two DSSO experiments, a total of 700 unique inter-linked peptides from 236 proteins were identified (**Fig. 1c** and **Supplementary Dataset 1**). 42.4% of these crosslinks mapped to Uniprot-annotated membrane-spanning proteins, demonstrating a strong focus on this compartment. The PhoX sample, processed using IMAC and SEC (see **Methods**), resulted in 85.3% of total peptides demonstrating a crosslinked lysine (669 unique inter-links, 1257 loop-links, 6534 uninformative mono-links), derived from 782 proteins (**Fig. 1d** and **Supplementary Dataset 2**). While enrichment for membrane-spanning proteins for PhoX was less than DSSO, at 27.9 %, this value was still broadly consistent with our prior studies using CSC alone^7^. Combining DSSO and PhoX data, our “structural surfaceomics” approach identified 2,390 total inter-linked and intra-linked peptides on Nomo-1 cells.

**Figure 2.**
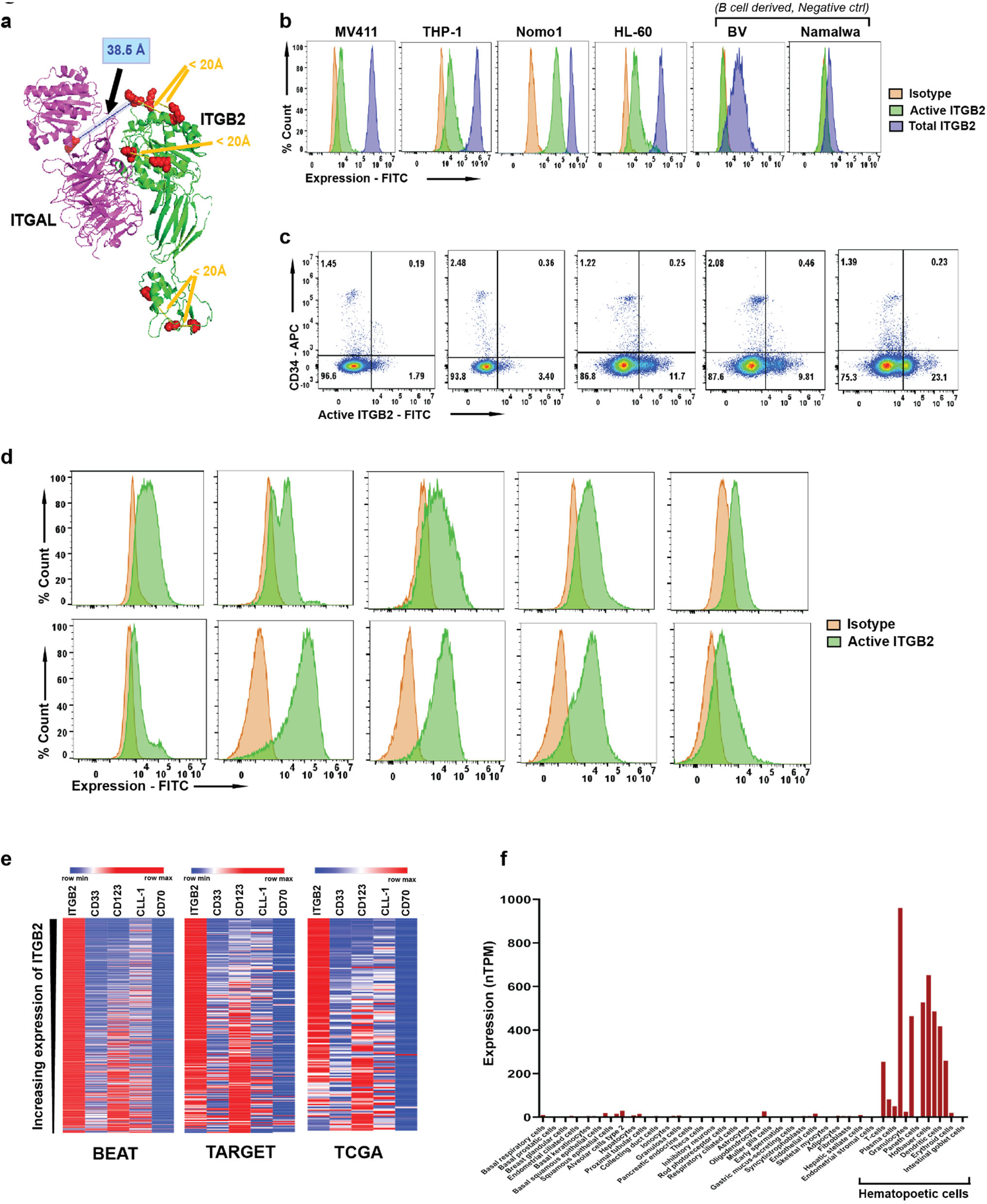
Activated Integrin-β2 is conformationally selective antigen in AML. **a.** Identified cross-linked peptides mapped on to the crystal structure of integrin-αL/integrin-β2 heterodimer (PDB: 5E6R). **b.** Flow cytometry histogram plot showing expression of total and activated integrin-β2 on AML and B-cell lines (BV and Namlwa). The y-axis represents percent count normalized to mode. Gating strategy shown in (Extended Data Fig. 3b). Representative plots from *n* = 3 independent experiments. **c.** Flow cytometry plot showing absence of active Integrin-β2 on CD34+ HSPCs from GM-CSF mobilized peripheral blood. Gating strategy shown in (Extended Data Fig. 3d). Deidentified patient samples were used for this analysis (*n* = 5 independent donors). Representative of 1-2 independent experiments. **d.** Representative flow cytometry histogram plots showing expression of active Integrin-β2 on primary AML cells. The y-axis represents percent count normalized to mode. Gating strategy shown in (Extended Data Fig. 3e). (Representative of *n* = 10 total deidentified samples, performed in single assay each). **e.** Heat map showing inverse expression pattern of *ITGB2* against other AML targets in publicly available primary AML RNA-seq data. Color bar represents maximum expression in each row based on normalized read counts. Sample size of BEAT^40^ AML (adult), TARGET^41^ (pediatric) and TCGA^39^ were 510, 255 and 150 respectively. **f.** Aggregated single cell RNA-seq data showing essentially exclusive expression of *ITGB2* in hematopoietic tissue, obtained from the Human Protein Atlas^42^.

### Active integrin-β2 as a potential conformation-selective target in AML

We manually inspected the crosslinked peptides obtained from our structural surfaceomics analysis, with our primary metric being comparison to published structures in the Protein Data Bank. In our DSSO data, we were particularly intrigued to find several crosslinks mapping to the protein integrin-β2 as well as its heterodimer partner integrin-α_L_ (PDB:5E6R)^28^. We first noted several intra-protein cross-links within integrin-β2 itself that fell within the C_α_ Lys-Lys distance constraints of the DSSO cross-linker, < 20 Å. However, we found four cross-links that did not match the C_α_-C_α_ distance constraint on the available crystal structure, extending to ∼38.5 Å between Lys194 and Lys196 of the βI domain of integrin-β2 and Lys305 and Lys330 on the I domain of integrin-α_L_ (ref.^28, 29^) (**Fig. 2a**). Notably, the crystal structure appears to represent the inactive, closed form of this integrin heterodimer^28, 29^. Our XL-MS data suggested that these domains are instead in closer proximity on Nomo-1, potentially consistent with the open, active conformation in these AML tumor cells (**Extended Data Fig. 3a**).

This finding was notable as integrin-β2 has been identified on several immune cell types including monocytes, neutrophils, NK cells, and T cells^30, 31^. However, at the protein level, it is known to largely remain in the closed, resting conformation until cellular activation, after exposure to appropriate cytokines, adhesion molecules, or other proteins^32–35^. Furthermore, a previous study suggested that constitutive signaling through integrin-β2 maintains proliferation in AML blasts^36^. Taken together, these results suggest that aberrant AML biology may lead to constitutive activation of integrin-β2, thus creating a possible tumor-specific conformation that, when targeted, would largely spare normal, resting hematopoietic cells.

**Figure 3.**
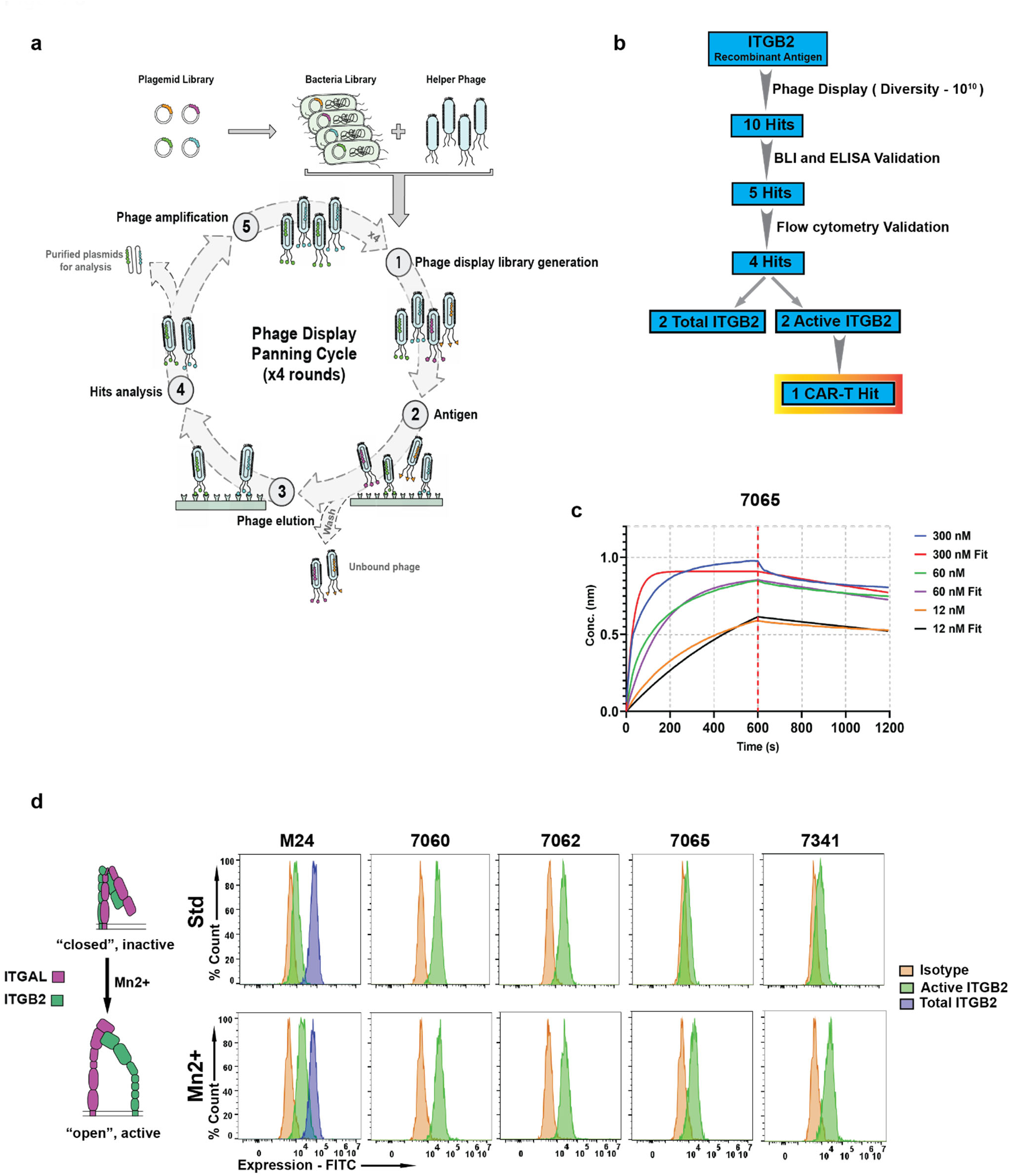
Antibody 7065 binds preferentially to the active conformation of Integrin-β2. **a.** Schematic flow diagram of phage display selection strategy used for developing anti-Integrin-β2 antibodies. **b.** Schematic flow diagram showing triage of antibodies obtained from phage display library and the downstream validation/funneling to identify an active integrin-β2 binder. **c.** Representative biolayer interferometry plot showing determination of binding affinity (KD) of 7065 antibody against integrin-αL/integrin β2. *n* = 3 different concentrations of antibody was used for this experiment (also see Extended Data Fig. 6c). **d.** Flow cytometry analysis of Jurkat T-ALL cells in presence and absence of 2 mM Mn^2+^ ions, to determine/identify antibodies having specificity against active integrin-β2. The y-axis represents percent count normalized to mode. Gating strategy shown in (Extended Data Fig. 3b). (representative of *n* = 2 independent experiments)

To explore this hypothesis, we took advantage of the murine monoclonal antibody “M24”, widely used to selectively recognize the activated form of integrin-β2 by flow cytometry^37^. We profiled four AML cell lines of varying genotype (Nomo-1, THP-1, HL-60, MV4-11) and confirmed that all showed clear expression of activated integrin-β2 by M24 staining, in addition to high levels of total integrin-β2 by TS1/18 clone (**Fig. 2b**). In contrast, B-cell malignancy lines BV and Namalwa showed total integrin-β2 but no discernable activated conformation expression (**Fig. 2b**). To extend this result to normal hematopoietic progenitors, we further obtained GM-CSF mobilized peripheral blood samples from five hematopoietic stem cell transplant donors at our institution. We found that CD34+ hematopoietic stem and progenitor cells (HSPCs) from these individuals showed no evidence of activated integrin-β2 by flow cytometry (**Fig. 2c**), though they did express total integrin-β2 (**Extended Data Fig. 3c**). This result provides an initial suggestion of a favorable therapeutic index for this target.

To further evaluate activated integrin-β2 in primary AML, we obtained de-identified bone marrow aspirate specimens from ten patients at our institution (**Fig. 2d**) and two patient derived-xenograft (PDX) models of AML from the PRoXe biobank^38^ (**Extended Data Fig. 3f**). Gating on the mature blast population, we found that activated integrin-β2 appeared highly expressed in nine of twelve total samples analyzed. We further analyzed bulk RNA-seq data across three AML patient tumor datasets (TCGA and BEAT AML: adult; TARGET: pediatric)^39–41^, finding high levels of expression of *ITGB2* transcript across patient blasts (**Fig. 2e**).

Interestingly, we found a complimentary expression pattern of *ITGB2* with leading AML targets *CD33* and *IL3RA* (CD123), suggesting that tumors with low expression of these current leading antigens may potentially benefit from anti-integrin-β2 therapy (**Fig. 2e**). We also found consistent, high expression of *ITGB2* across various AML genotypes (**Extended Data Fig. 4a**). However, we do note that transcript expression alone cannot report as to whether surface integrin-β2 is in the activated or resting conformation. Toward the safety profile of this target, we evaluated aggregated single cell RNA-seq data in the Human Protein Atlas^42^. We noted that *ITGB2* transcript is only detectably expressed on hematopoietic cell types (**Fig. 2f**), with high expression across the myeloid lineage^34, 35^. Already, this transcript expression pattern compares favorably with that of other known AML immunotherapy targets (**Extended Data Fig. 4b**). However, we anticipate that conformation-selective targeting will lead to an additional layer of discrimination between tumor and normal cells not available to these other targets.

**Figure 4.**
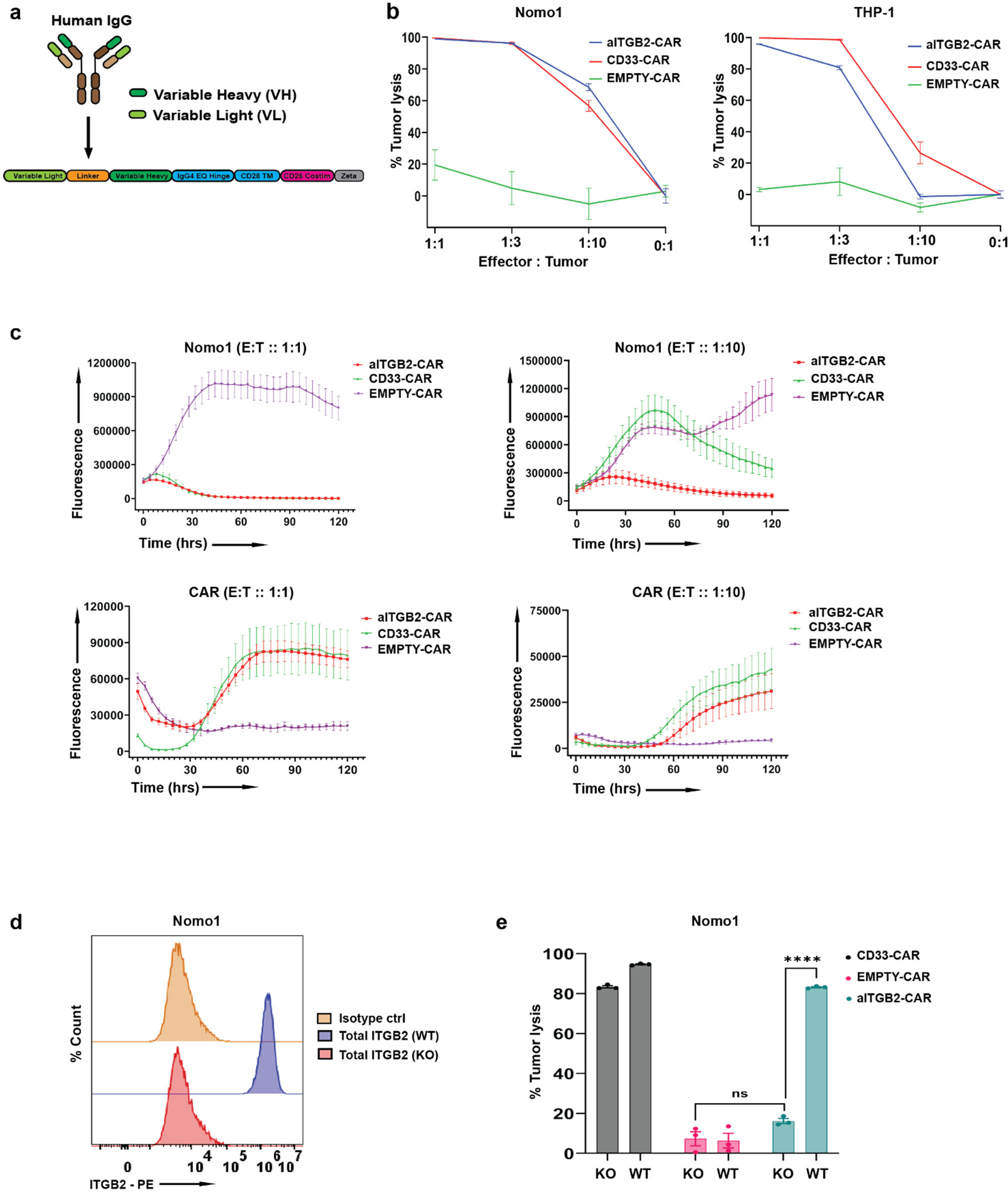
Anti-active integrin-β2 (aITGB2) CAR-T derived from 7065 antibody is cytotoxic to AML cells. **a.** Schematic diagram of CAR-T construct used. **b.** Luciferase based cytotoxicity of aITGB2 CAR-T design against Nomo1 and THP-1 AML cell lines. *n* = 3 technical replicates, representative plot from 4 independent experiments. **c.** Incucyte live-cell imaging data demonstrating efficient cytotoxicity of aITGB2 CAR-T against Nomo1 at 2 different E:T ratio, 1:1 and 1:10 over 5 days period. CAR-T cells were labelled with GFP and tumor cells (Nomo-1) with mCherry to facilitate fluorescence-based quantification. The *y*-axis represents integrated fluorescence used as a proxy to monitor cell proliferation. Performed with *n* = 6 technical replicates. **d.** Flow cytometry histogram showing successfully generated *ITGB2* knockout version of Nomo-1 using CRISPR-Cas9. The y-axis represents percent count normalized to mode. Gating strategy shown in (Extended Data Fig. 3b). Representative of *n* = 3 independent experiments. **e.** Luciferase based cytotoxicity data showing specific activity of aITGB2 CAR-T against WT Nomo-1 and not against its *ITGB2* knockout Nomo-1 (E:T ratio was 1:1 with overnight incubation). *n* = 3 technical replicates. The luciferase signals of the cytotoxicity assays in this figure were normalized against untransduced CAR-T of their respective E:T ratios. All statistical data in this figure are represented as mean ± SEM, with *p*-value by two-tailed *t*-test.

### Characterization of recombinant antibody binders versus active integrin-β2

Our next goal was to develop chimeric antigen receptor (CAR) T cells versus active integrin-β2 as a proof-of-principle therapeutic for AML. We first explored two commercially available antibody clones versus active integrin-β2, M24 (ref.^43^) and AL57 (ref.^44^). Using the sequence of these antibodies, we designed single chain variable fragment (scFv) binders and incorporated them into a CD28-based CAR backbone. While we found no activity for AL57-based scFv’s, we did find that both the designs (V_H_-V_L_ and V_L_-V_H_) of the M24-derived scFv did indeed lead to some Nomo-1 cytotoxicity (**Extended Data Fig. 5**). Here and throughout the study, we also used a previously-described anti-CD33 CAR as a positive control^20^.

**Figure 5.**
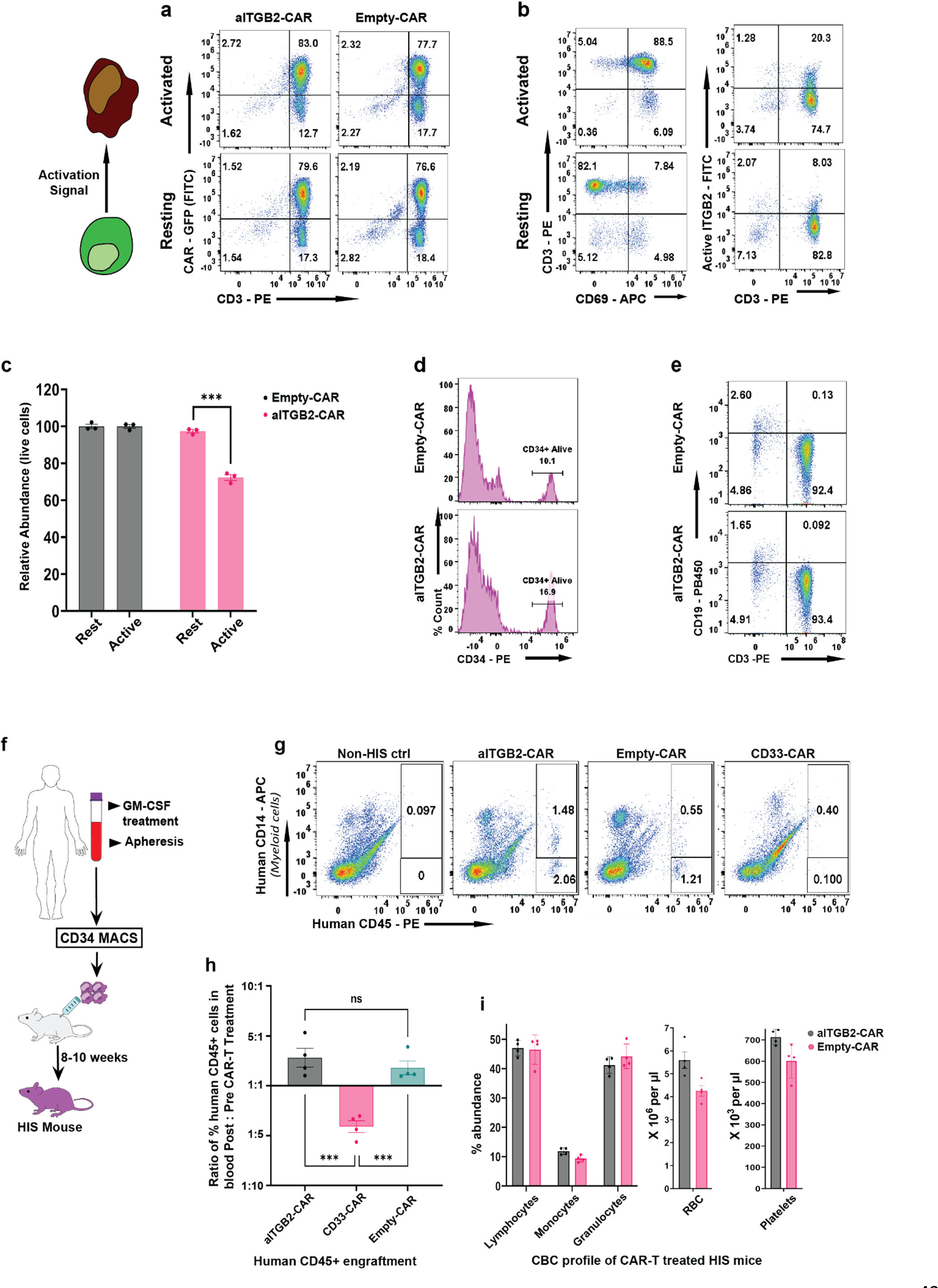
Toxicity assessment of aITGB2 CAR-T demonstrates a promising safety profile. **a.** Representative flow cytometry-based cytotoxicity assay showing specificity of aITGB2 CAR-T against activated peripheral blood T cells which harbors activated integrin-β2 (focus on lower right quadrant, with CAR-negative, CD3-positive T-cells). Both resting and activated conditions performed in overnight co-culture assays with aITGB2 CAR-T cells. (Gating strategy similar to shown Extended Data Fig. 3d.) **b.** Representative flow cytometry analysis showing successful activation of T cells and partial abundance of activated integrin-β2 on activated T cells. (Gating strategy similar to shown in Extended Data Fig. 3d.) **c.** Quantitative analysis of active T-cell depletion data in (a); *n* = 3 technical replicates. **d.** Representative flow cytometry analysis showing no discernible impact of aITGB2 CAR-T against CD34+ HSPCs from GM-CSF mobilized peripheral blood. The y-axis represents percent count normalized to mode. (Gating strategy similar to that shown in Extended Data Fig. 8c) (n = 1 donor) and similarly for **e.** T cells and B cells. (Gating strategy similar to shown in Extended Data Fig. 3d.) (n = 3 technical replicates and representative of 2 independent experiments). Also see Extended Data Fig. 9c. **f.** Schematic flow diagram for generation of humanized immune system (HIS) mice. **g.** Representative flow cytometry data from HIS mice data showing apparent non-toxicity of aITGB2 CAR-T against myeloid cells (CD14+). All events were used for gating and analysis. (Representative plot from n = 4 – 6 mice and 6 days post CAR-T treatment). **h.** Quantification of hCD45+ data in (g). Gating strategy similar to shown (Extended Data Fig. 10d). *p*-value by two-tail *t*-test. \*\*\**p* < 0.005. **i.** Complete blood count profiling of HIS mice treated with aITGB2 CAR-T at day 5 (data from *n* = 4 mice). All the statistical data in this figure are represented as mean ± SEM. For all the *in vitro* cytotoxicity assays, E:T ratio was 1:1 with overnight incubation time.

While this result was promising that CAR-T’s could be developed versus active integrin-β2, these M24-derived CAR-T’s showed relatively limited *in vitro* potency versus Nomo-1when compared to anti-CD33 CAR-T. Furthermore, the M24 framework sequences are fully murine^43^, increasing potential for immunogenicity when used in a human therapeutic. Therefore, we sought to develop alternative CAR-T cell designs.

As a first step, we used our previously-described Fab-phage display platform^45^, based on a fully human framework sequence, to perform selections versus recombinant integrin-β2 (**Fig. 3a** and **Methods**). From the initial library diversity of ∼10^10^ binders, we identified ten initial hits versus integrin-β2, five of which were validated using bio-layer interferometry (BLI) and non-specific ELISA (**Fig. 3b, c** and **Extended Data Fig. 6**) to have binding affinities to integrin-β2 in the low-nM range and lack of binding to irrelevant proteins, respectively (**Extended Data Fig. 6b, c** and **Supplementary Table 1**). These five Fabs were cloned into a human IgG1 backbone and were purified following recombinant expression in mammalian cells (**Extended Data Fig. 6**). As a validation system, we chose the Jurkat T-ALL cell line, which we found expresses high levels of integrin-β2 with a fraction appearing to show constitutive activation at baseline based on M24 staining (**Fig. 3d**). Encouragingly, we found that four of our five recombinant antibodies versus integrin-β2 showed positive signal by flow cytometry on Jurkat (**Fig. 3d**).

**Figure 6.**
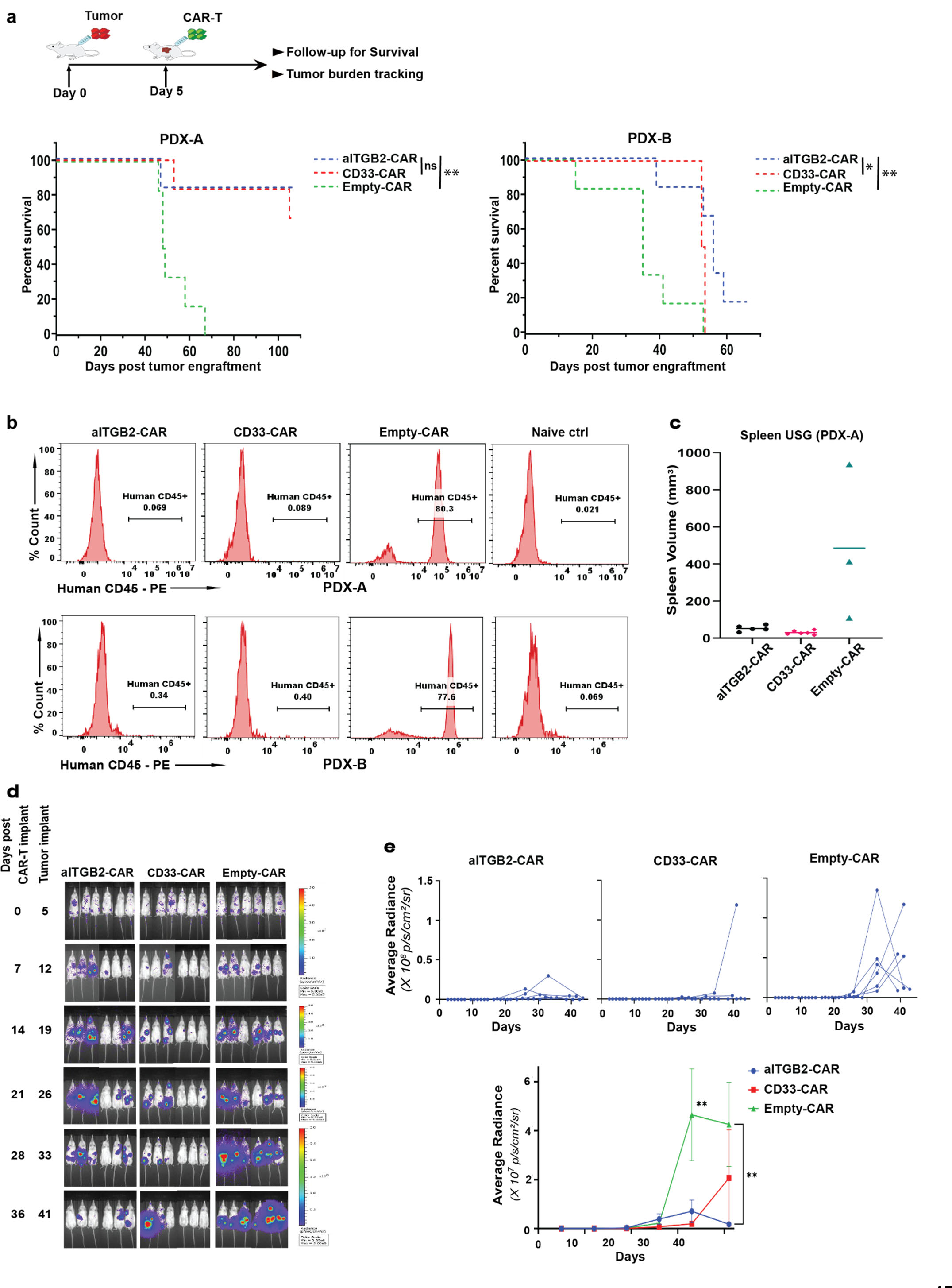
Efficacy of aITGB2 CAR-T against AML models *in vivo*. **a.** Survival of NSG mice implanted with 2 independent AML PDX and treated with aITGB2, anti-CD33, or empty CAR-T cells. *n* = 6 mice per arm. *p*-value by log-rank test. 2 million AML tumor cells injected on Day 0, 5 million CAR-T cells injected on Day 5. **b.** Representative flow cytometry histogram plots of peripheral blood draw showing tumor burden at 8-week post tumor injection for PDX-A and 3.5 weeks for PDX-B. (also see Extended Data Figure 10a, b.). Naïve control mice have no human cells (AML tumor or CAR-T) injected and used to assess background noise in flow cytometry assay. The *y*-axis represents percent count normalized to mode. Gating strategy similar to shown (Extended Data Fig. 10d). Representative of data from *n* = 4 - 6 mice per arm dependent on number of mice alive until that time point. **c.** Spleen ultrasonography from Empty CAR-treated group compared to CD33 or aITGB2 CAR-T treated mice. All mice alive at day 49 post tumor implantation were scanned (*n* = 3 - 6 mice/arm still surviving at this time). **d.** BLI imaging showing efficacy of aITGB2 CAR-T against intravenously implanted AML cell line Nomo-1 (*n* = 6 mice/arm). **e.** Quantitative analysis of bioluminescence intensity of these mice plotted individually (*n* = 6). Mann Whitney test was used for statistical analysis of mice bioluminescence quantification. All the statistical data in this figure are represented as mean ±SEM.

We next took advantage of the fact that integrins can be biochemically converted from the inactive, closed conformation to the active, open conformation by treatment with the divalent cation Mn^2+^ (ref.^46^). While two clones (7060, 7062) did not show any responsiveness to 2 mM Mn^2+^ treatment, clones 7065 and 7341 showed increased signal in response to Mn^2+^ (**Fig. 3d**). Indeed, the profile of clone 7065 appeared highly similar to that of the well-validated antibody M24, with limited signal in the absence of Mn^2+^ but ∼3-fold increased median fluorescence intensity after cation exposure. The higher signal from 7341 at baseline suggests that it may also have some binding to the closed conformation of integrin-β2, in addition to recognizing the active conformation. These findings suggest that clone 7065 may be particularly selective for the activated conformation of integrin-β2.

### Development of anti-active integrin-β2 CAR-T cells

Toward CAR-T generation, the sequences of clones 7065 and 7341 were engineered into scFv format and cloned into a backbone with a CD28 co-stimulatory domain (**Fig. 4a**). For each antibody we again tried two different scFv orientations, either V_H_-V_L_ or V_L_-V_H_ with a 3x Gly_4_Ser linker. Based on Nomo-1 cytotoxicity *in vitro*, the 7065 V_L_-V_H_ design appeared to be most efficacious (**Extended Data Fig. 7a**) compared to control “empty” CAR-T cells (construct with full CAR backbone but no antibody binder). This 7065 design also showed no discernible activity versus a negative control of AMO-1 multiple myeloma cells, which do not express activated integrin-β2 (**Extended Data Fig. 7a, c**).

While these initial *in vitro* experiments were promising, we did anecdotally observe decreased proliferation and final yield of these CAR-T cells during manufacturing, compared to other CAR-Ts produced in our group. We also noted that even the best performing CAR-T design had moderate Nomo-1 cytotoxicity compared to the positive control anti-CD33 CAR-T (**Extended Data Fig. 7a**). We hypothesized that T-cell stimulation was leading to integrin-β2 activation, and thus some degree of CAR-T “fratricide” during expansion, eliminating some CAR-Ts and negatively impacting others due to constant activation. To test this hypothesis, we used an approach employed in manufacturing for other CAR-T targets present on activated T-cells, such as CD70 (ref.^47^). Namely, we used a CRISPR-Cas9 ribonucleoprotein (RNP) strategy to knock out *ITGB2* prior to T-cell stimulation and lentiviral transduction. We evaluated four sgRNA designs and found sgRNA-1 and 4 showed high knockout efficiency (**Extended Data Fig. 7d**). Using this manufacturing protocol, we no longer observed any deficit in CAR-T expansion (**Extended Data Fig. 7e**), and, furthermore, we observed *in vitro* cytotoxicity versus Nomo-1 and THP-1 cells comparable to anti-CD33 CAR-T (**Fig. 4b**). The CAR-Ts were also found to have potent degranulation against Nomo-1 (**Extended Data Fig. 8a**).

We further varied the V_L_-V_H_ linker length between 1x-4x Gly_4_Ser and found largely consistent cytotoxicity (**Extended Data Fig. 8b**). In assays below, we thus chose either the 3x or 4x linker designs as lead candidates for further evaluation. To assess proliferation kinetics of these anti-active integrin-β2 CAR-T (aITGB2) designs, we performed live cell imaging assays of Nomo-1 co-culture. We found that at 1:1 Effector to Tumor (E:T) ratio, aITGB2 CAR-Ts showed similar proliferation and cytotoxicity to anti-CD33 CAR-Ts (**Fig. 4c**). However, at 1:10 E:T, aITGB2 CAR-Ts outperformed anti-CD33 CAR-T (**Fig. 4c**). Both CAR-Ts showed similar proliferation in this co-culture assay (**Fig. 4c**). Profiling of aITGB2 and CD33 CAR-T pre- and post-tumor exposure demonstrated similar expression of memory-like phenotype markers based on CD62L and CD45RA staining (**Extended Data Fig. 8d**). Taken together, these findings encourage further preclinical investigation of our aITGB2 CAR-Ts as an AML therapy.

### aITGB2 CAR-T is specific against the active conformation of integrin-β2

We next evaluated specificity of our CAR-T cell for the active conformation of integrin-β2. First, we used our Cas9 RNP strategy to confirm that *ITGB2* knockout in Nomo-1 fully abrogated aITGB2 CAR-T activity (**Fig. 4d, e**). While this finding supports that our CAR-T is specific to integrin-β2, it does not confirm conformation specificity. As a first test, we confirmed no aITGB2 CAR-T cytotoxicity versus the B-cell leukemia line Namalwa, which expresses total integrin-β2 but not the active conformation based on M24 staining (**Extended Data Fig. 9a**, **Fig. 2b**). As a second test, in an overnight assay we incubated GFP-labeled aITGB2 CAR-Ts with normal donor peripheral blood mononuclear cells (PBMCs). At baseline, we found that aITGB2 showed no cytotoxicity versus resting CD3+ T-cells, which are positive for total integrin-β2 but not the activated conformation (**Fig. 5a, e** and **Extended Data Fig. 9b**). However, with PBMC stimulation using ionomycin, lipopolysaccharide and IL-2 overnight during aITGB2 CAR-T co-culture, we found that there was partial depletion of the GFP-negative (i.e. non-CAR-T, derived from PBMC) T-cell population (**Fig. 5a, c**). Indeed, this partial depletion was consistent with the fraction of T-cells we found to express active integrin-β2 after stimulation, which notably was a much smaller fraction than CD69-positive cells (**Fig. 5b**). These results suggest that aITGB2 CAR-T cells specifically eliminate target cells displaying the activated conformation of this protein, while ignoring cells expressing even high levels of total integrin-β2 in the inactive, closed conformation.

### aITGB2 CAR-T appears to have minimal toxicity versus normal hematopoietic cells

Given that *ITGB2* only appears expressed in hematopoietic cells (**Fig. 2f**), we focused our further toxicity analysis on these populations. By M24 flow cytometry on peripheral blood we showed that resting T- and B-cells did not express active integrin-β2 (**Extended Data Fig. 9b**). Analyzing granulocytes and monocytes, we did find that these cells appeared strongly positive for active integrin-β2; however, it is well known that this finding is an artifact of *ex vivo* activation of these cells after blood collection^48^. We reasoned that evaluating potential aITGB2 CAR-T cytotoxicity impacts versus myeloid cells would require *in vivo* studies, in the absence of this activation artifact.

However, prior to these *in vivo* studies, we first performed overnight *in vitro* co-culture assays of aITGB2 with GM-CSF mobilized peripheral blood. Consistent with lack of active integrin-β2 on CD34+ HSPCs by flow cytometry (**Fig. 2c**), we found no depletion of HSPCs after aITGB2 co-culture (**Fig. 5d**). Similarly, in PBMCs we observed no depletion of T-cells (**Fig. 5e**), consistent with our findings in **Fig. 5a**. Surprisingly, we saw a modest depletion of CD19+ B-cells compared to “empty” control; the mechanism for this effect is unclear, but it does not appear to be specific to aITGB2 CAR-T given similar depletion in anti-CD33 CAR-T (**Extended Data Fig. 9c**). As expected, based on known artifactual integrin-β2 activation (**Extended Data Fig. 9d**), and confirming *in vitro* potency of aITGB2 CAR-T versus primary cells, we found strong depletion of monocytes and neutrophils (**Extended Data Fig. 9e**).

We next moved into a “humanized immune system” (HIS) murine model, where CD34+ HSPCs isolated from GM-CSF mobilized peripheral blood are intravenously implanted into busulfan treated NSG-SGM3 mice^49^ (**Fig. 5f**). Mice were monitored by peripheral blood draw at 8 weeks post-implant to confirm hematopoietic engraftment, assessed by at least 1.5% circulating human CD45+ mononuclear cells. At this time, we treated all successfully engrafted mice (16 of 25 total implanted) with aITGB2, anti-CD33, or empty CAR-T cells and 6 days later sacrificed mice and analyzed peripheral blood. While rigorous quantification of CD14+ cells was not possible due to high variability in myeloid engraftment at the time of CAR-T treatment, we found no discernible depletion after aITGB2 CAR-T (**Fig. 5g**). Importantly, we found a significant depletion of total human CD45+ cells in PBMC obtained from blood draw after treatment with CD33 CAR-T (**Fig. 5h**). This result recapitulated expected toxicity of targeting this marker expressed on HSPCs and myeloid cells, and served as a positive control that the chosen time point is effective in discerning CAR-T impacts on normal human blood cells. In contrast, human CD45+ cells continued to expand in mice treated with either aITGB2 or “empty” CAR-Ts (**Fig. 5h**).

Furthermore, we probed the 7065 antibody clone and found it was cross-reactive with murine activated integrin-β2 (**Extended Data Fig. 9g**). This cross-reactivity gave us the opportunity to evaluate toxicity directly to murine hematopoietic cells. We thus performed complete blood count (CBC) analysis of murine peripheral blood from our HIS mouse study above. At 5 days after aITGB2 CAR-T treatment, we found no depletion of any murine PBMC types (**Fig. 5i**). Taken together, these results suggest that treatment with aITGB2 CAR-T may carry minimal toxicities to bystander immune cells, unlike CD33 CAR-T, thus underscoring a promising safety profile.

### aITGB2 CAR-T is efficacious against AML patient-derived xenografts (PDX) *in vivo*

Finally, we evaluated *in vivo* efficacy of aITGB2 CAR-T. We established 2 separate monocytic leukemia PDX obtained from PRoXe^38^, one from a female and the other from a male patient, via intravenous implantation in NSG mice. Both of these samples appeared to express active integrin-β2 based on M24 flow cytometry (**Extended Data Fig. 3f**). 5 days post implantation of 2 million PDX AML cells, we treated mice with 5 million empty, aITGB2, or CD33 CAR-Ts. Tumor burden was monitored by periodic peripheral blood draw, evaluating for human CD45+ mononuclear cells, and/or ultrasonography for spleen size (**Fig. 6b, c, Extended Data Fig. 10a, b**). Notably, in both of these PDX models we saw marked elimination of human CD45+ cells, as well as decreased spleen size, in aITGB2 or CD33 CAR-T treated mice, with prominent outgrowth of tumor cells in “empty” CAR control (**Fig. 6b, c**). In both models, survival was significantly improved in aITGB2 CAR-T-treated mice compared to empty control, and was similar between aITGB2 CAR-T and anti-CD33 CAR-T (**Fig. 6a**). We further evaluated anti-tumor efficacy of aITGB2 CAR-T in a Nomo-1 cell line xenograft mouse model implanted in NSG mice (**Fig. 6d, e**). Tumor burden was monitored non-invasively via stable luciferase expression. In this study we again noted improved tumor control over empty CAR-T, as well as similar efficacy of aITGB2 CAR-T and anti-CD33 CAR-T (**Fig. 6d, e**). However, in this aggressive model, neither tested CAR-T could lead to complete tumor eradication. Toward initial investigation of a possible mechanism of relapse after aITGB2 CAR-T, we performed flow cytometry on murine spleens harvested after sacrifice at Day 42 post-tumor implant. Gating on human CD45+ AML blasts, we found no evidence of tumor downregulation or loss of activated integrin-β2 (**Extended Data Fig. 10c**). This initial experiment suggests that loss of the activated conformation of ITGB2 may not be an immediate mechanism of resistance to our structurally-selective targeting.

## DISCUSSION

Our structural surfaceomics approach presented here, integrating XL-MS with cell surface glycoprotein enrichment, is a technology designed to expand the targetable space of cell surface immunotherapy antigens. Using this strategy, we identified the active, open conformation of integrin-β2 as a promising immunotherapy target in AML, a hematologic malignancy in significant need of new therapeutic options. We further developed humanized scFv-based CAR-T cells against active integrin-β2 and found them to be both safe and efficacious using *in vitro* and *in vivo* models. Taken together, our results demonstrate a first application of a potential pipeline for conformation-selective immunotherapy target discovery, not possible with traditional transcriptome- or proteome-focused abundance analysis.

We believe that the structural surfaceomics approach carries promise in applications not only for immunotherapy target discovery, but also basic or translational science in other fields. These could range from infectious disease to neuroscience, where obtaining low-resolution structural information on a broad swath of plasma membrane proteins may spur new areas of investigation. However, we do acknowledge that our current structural surfaceomics approach carries limitations. First, sample input: XL-MS has traditionally required large sample inputs (10^9^ cell scale) and extensive mass spectrometer time to identify cross-linked peptides. These limitations led us to focus our initial efforts here on a single AML cell line with multiple XL-MS approaches. However, future optimization of enrichable cross-linkers, alternative cross-linker reactivities, as well as further technological MS advances, may enable broader scale profiling of both tumor and normal cells, or even primary samples. Second, analysis and validation of potential targets: in the current study we manually compared identified crosslinks to PDB structures to find targets of interest. Future work will aim to develop automated computational structural analysis to identify the most promising targets for workup. In terms of validation, we chose to first investigate integrin-β2 in depth because we had flow cytometry and biochemical (i.e. Mn^2+^) tools by which to probe its conformation status. For other potential targets these tools will not exist *a priori*. We thus anticipate future efforts to develop alternative strategies (for example, “disulfide locking”, as used in many structural biology studies of membrane proteins^50^) to generate putative tumor-selective conformations for recombinant antibody selection and subsequent validation.

The active conformation of integrin-β2 carries particular promise compared to other known AML immunotherapy targets given a potentially improved safety profile, with no discernible activity versus HSPCs or resting myeloid cells. While we do anticipate there will be some unwanted activity versus activated myeloid or T-cells, we predict this toxicity will still be significantly lower than other AML targets such as CD33, CD123, or CLL-1 that are expressed widely on all mature myeloid cells^51^. Our results also suggest that depletion of activated T-cells may be limited in humans, as *in vitro* only a fraction of donor T-cells appeared to express active integrin-β2 even after potent stimulation.

In terms of efficacy, like many other AML targets^16, 52^, we observed heterogeneity of active integrin-β2 on primary patient tumor samples. Therefore, we acknowledge that aITGB2 CAR-T is unlikely to be a curative therapy for all AML patients. However, for tumors with elevated expression of this target, in our *in vitro* and *in vivo* experiments we did not observe distinctly decreased efficacy versus anti-CD33 CAR-T, a leading AML CAR-T target but with marked toxicity concerns^19^. The favorable safety profile of aITGB2 CAR-Ts also may create future opportunities for multi-targeting CARs versus two or more antigens with complementary but heterogeneous tumor expression patterns, particularly if the additional antigens beyond active integrin-β2 also are non-myeloablative. Future antibody engineering efforts, or incorporation of recently-described chimeric CAR-TCR designs^47^, may be able to enhance efficacy of aITGB2 CAR-Ts versus tumor cells expressing low antigen levels.

In conclusion, our studies demonstrate a potential systematic approach to identify and target conformation-specific antigens in cancer. Humanized aITGB2 CAR-Ts, discovered via this approach, stand as a promising proof of principle therapeutic warranting further preclinical evaluation in AML and a pathway for many other applications of structurally directed immunotherapeutic targets.

## Author Contributions

K.M. and A.P.W. conceptualized the study, acquired the funding, performed data analysis/interpretation, and wrote the manuscript. K.M., G.W., C.Y., J.J.A., W.C.T., B.P.E., M.G., M.R.H., C.H.I., A.N., J.A.C.S., F.S., P.P. and B.J.H. performed experiments and/or data analysis. E.R., C.K., M.M., E.S. and C.C.S.: primary patient sample acquisition and/or analysis. A.P.W., M.A.N., V.S., A.S., S.S.S., L.H. and R.L.M. provided resources and/or supervised the study.

## Acknowledgements

We thank Prof. Dean Sheppard (UCSF) for providing his expert opinion and consultation on integrin biology. We also thank Dr. Susanna K. Elledge (UCSF) and Dr. Mark A. Burlingame (UCSF) for technical assistance with MS sample analysis. We thank Neil Wiita for assistance in figure graphics. We also thank patients and their families who contributed research specimens to associated tissue banks. We acknowledge funding from the Michelson Prize-2019 (to K.M.) awarded by Michelson Medical Research Foundation and Human Vaccine Project; NIH R21 CA263299 (to A.P.W.); NIH R01GM074830 and NIH R01GM130144 (to L.H); Canadian Institutes of Health Research (MOPS-136944) and from Bristol-Myers Squibb (to S.S.S); National Cancer Institute Cancer Center Support Grant P30CA082103 (to the University of California, San Francisco) to support the Pediatric Hematopoietic Tissue Cell Bank); NIH R01GM087221, NIH S10OD026936, and the National Science Foundation award 1920268 (to R.L.M and M.R.H); American Society of Hematology Research Training Award for Fellows and Chan Zuckerberg Biohub Physician-Scientist Fellowship Program (to W.C.T). Flow cytometry was performed at the UCSF Laboratory for Cell Analysis and murine studies performed at the UCSF Preclinical Therapeutics Core, both part of the Helen Diller Family Comprehensive Cancer Center and supported by P30 CA082103.

## Conflicts of Interest

K.M., J.J.A., S.S.S., and A.P.W. have filed a provisional patent related to the antibody sequences described herein. A.P.W. has received research funding from Genentech. C.S. has received research funding from Revolution Medicines, Abbvie and Erasca, Inc. and has served on advisory boards for Genentech, Abbvie and Astellas. All other authors declare no conflict of interest.

## Materials and Methods

### Cell lines, PDX and patient samples

Nomo-1 cell line was obtained from DMSZ. THP1, HL60, MV411, Jurkat and S49.1 were obtained from ATCC. All cell lines were grown in RPMI-1640 media (Gibco, 11875093) with 20% FBS (BenchMark, Gemini, 100-106) and 100 U/ml Penicillin-Streptomycin (UCSF Cell Culture Facility). All the cells were grown in 5% CO_2_ at 37° C. All AML PDX were procured from Public Repository for Xenografts (PRoXe) at Dana-Farber Cancer Center under an appropriate Materials Transfer Agreement. Primary AML samples were obtained from the UCSF Hematologic Malignancies Tissue Bank and the Pediatric Hematopoietic Tissue Cell Bank under protocols approved by the UCSF Committee on Human Research Institutional Review Board (IRB).

### Cross-linking and Cell surface labelling

The DSSO (Sigma Aldrich, 909602) based XL-MS involving high-pH fractionation, and PhoX (Thermo Fisher Scientic, A52286) based XL-MS was each performed with 2.4 X 10^9^ cells (in batches of 6 X 10^8^). However, the initial DSSO experiment without high-pH fractionation was done with 4 X 10^8^ cells. For each experiment, the cells were harvested and washed (300 RCF for 5 min) thrice with PBS each time to get rid of all the amine containing components of the media and finally resuspended in PBS. Then the amine reactive cross-linker DSSO or PhoX pre-dissolved in DMSO (Sigma Aldrich, 276855) is added to the cells at a final concentration of 10mM and incubated at RT for 45 minutes. The cross-linking step was followed by biotinylation of the cell surface proteins using glycoxidation chemistry of the N-linked glycosylation-site.

Briefly, the cells were then washed with PBS thrice and treated with 1.6 mM sodium metaperiodate (VWR, 13798-22) for 20 minutes at 4C for oxidation of the N-linked sugar residues. The cells were again washed twice with PBS and treated with 10 mM aniline (Sigma-Aldrich, 242284) and 1 mM biocytin hydrazide (Biotium, 90060) for 90 minutes at 4° C, for installation of biotin on the oxidized sugar residues. The cells were then washed thrice to get rid of the excess of biotinylating reagents and snap froze in liquid nitrogen, and stored at -80° C until further processing. All the incubation steps were carried out in end-to-end rotor for gentle mixing during the reactions.

### Cell surface proteomics sample preparation

The frozen cell pellets were thawed in ice and were resuspended in 1 ml RIPA lysis buffer (Millipore Sigma, 20-188) with Halt protease inhibitor (Thermo Fisher Scientific, 78430) and 1 mM EDTA (Invitrogen, 15575-038). The cell suspension was then sonicated to lysis the cells followed by incubation in ice for 10 minutes with intermittent vortexing every 2-3 minutes. The lysate was then centrifuged at 17000 RCF for 10 minutes at 4o C to get the clarified supernatant containing the biotinylated cell surface proteins. This clear supernatant was added to the 0.5 ml of Neutravidin beads (Thermo Fisher Scientific, PI29204) prewashed and equilibrated with RIPA lysis buffer + 1 mM EDTA. This pulldown step was allowed to happen at 4° C for 2 hours. To remove non-specifically bound proteins, the beads were washed extensively using vacuum manifold (Promega), consecutively with 50 mL RIPA lysis buffer + 1mM EDTA, 50 mL PBS + 1 M NaCl and 50 mL 2 M Urea (VWR, 97063-798) + 50 mM Ammonium Bicarbonate. The beads bound with biotinylated cell surface proteins were resuspended in 50 mM Tris (pH 8.5) + 4 M urea + 10 mM TCEP (Gold Biotechnology, TCEP10) and 20 mm IAA (VWR, 97064-926). 10 ug Trypsin-LysC (Thermo Fisher Scientific, PRV5073) mix was added to this mixture to allow on-bead digestion of the bound proteins for simultaneous reduction and alkylation of cysteines residues at RT in end-to-end rotor. At 4 M urea, LysC continues digestion for 2 hours after which the mixture is diluted to 1.5 M urea using 50 mM tris (pH 8.5) upon which trypsin also gets activated and this protease digestion goes overnight (16-20 hours). The solution is centrifuged to pellet down the beads and the supernatant contained the tryptic peptides were transferred to fresh tube and acidified with 0.5% Trifluoroacetic acid (TFA). The peptides were then desalted using SOLA HRP Column (Thermo Scientific, 60109-001) and eluted with 50% acetonitrile (ACN) + 0.1% formic acid (FA). Finally, the peptides were dried down in speedvac (CentriVap, Labconco).

### Immobilized metal affinity chromatography (IMAC) purification for PhoX

Dry peptides were reconstituted in 80% ACN + 0.1% TFA. Meanwhile, Superflow Ni-NTA beads were stripped off using EDTA and reloaded with FeCl3 (Sigma Aldrich, 451649) on a polyprep chromatography column (Biorad, 7326008). Fe^3+^ loaded beads were transferred to C18 tips (Nest Group, SEM SS18V.25) where it was incubated for 4 - 6 minutes with intermittent mixing with the reconstituted peptides to allow specific binding of the PhoX (cross-linker with IMAC handle) bearing peptides. The beads were then rigorously washed with 0.5 % formic acid (FA) to rid of the unbound or the non-specifically bound peptides. The bound peptides were then eluted with 0.5 M Potassium Phosphate buffer (pH 7.4). The peptides eluted from the beads gets again gets bound to the C18 chromatographic material of the nest tips. The tips were washed thrice with 0.5 % FA and finally eluted with 50% ACN + 0.1 % FA and dried down in speedvac.

### Size-Exclusion Chromatography (SEC)

Size based fractionation of the peptides were done using Superdex Peptide 3.2/300 (GE Healthcare) column and HPLC (Agilent 1260 Infinity II). The dried peptides were reconstituted in the mobile phase constituting 30% ACN + 0.1% TFA and loaded on to the column. The run time was 90 minutes at a flow rate of 50 µl/min and 45 fractions (2 minutes per fraction) were collected in total. The fractions associated with the desired molecular weight were dried down in speedvac and stored at -80 for MS analysis.

### LC-MS and data analysis for data-dependent acquisition (DDA) proteomics

The peptide samples prepared for building Nomo-1 sufaceome custom database were loaded on to the an EASY-Spray nanocolumn (Thermo Fisher Scientific, ES900) installed on Dionex Ultimate 3000 NanoRSLC instrument coupled with Q-Exactive Plus mass spectrometer (Thermo Fisher Scientific). Peptides were separated over a 313 minute gradient of ACN ranging from 2.4% to 32% ACN and subsequently stepped up to 80% ACN over next 10 minutes, all with a flow rate of 0.3 μL/min. MS scans were performed over mass range of *m/z* 299-1799 with resolution of 70,000 FWHM at m/z 200. The resolution for MS/MS scans was set to 17,500 FWHM at *m/z* 200. Normalized collision energies of 27, 30 and 33 in stepped higher collision-induced dissociation (HCD) mode was used for fragmentation of the topmost 15 most intense precursor ions with isolation window of 1.7 *m/z*. To avoid the repeated sampling of high abundant ions, dynamic exclusion was turned on and set to 20 seconds. The data collected for MS and MS/MS was in profile mode centroided mode, respectively.

MS generated .raw files were processed using MSFragger53 within FragPipe with default settings unless stated otherwise. Briefly, the spectral data were searched against the human proteome database (UniProt, downloaded 05/11/21, 20,395 entries). The contaminant and decoy protein sequences were added to the search database using the inbuilt feature of the FragPipe pipeline downstream statistical analysis. The search was run with “Mass calibration and parameter optimization” and “closed search default config” allowing ±20 ppm mass tolerance for precursor ions and ±20 ppm for that of fragment ions. The inbuilt tools PeptideProphet and ProteinProphet were used for statistical validation of search results and subsequent mapping of the peptides to the proteins respectively with 1% FDR.

### High pH reverse-phase tip (HpHt) based fractionation of DSSO cross-linked peptides

The SEC fractions 13 and 14 which are enriched with DSSO cross-linked peptides (**Extended Data Fig. 1b**) were further fractionated by high pH reverse-phase tip (HpHt) as described previously^27^. Briefly, the HpH tip was constructed in a 200-µL pipette tip by packing C8 membrane (Empore 3M) and 5 mg of C18 solid phase (3 μm, Durashell, Phenomenex). The HpHt column was sequentially washed with a series of 3 different solvents/solutions namely methanol, ACN and ammonia water (pH 10), 90 µl each. Then, each SEC fraction was loaded onto the HpHt column, which was centrifuged at 1,200 PRM for 5 min. The bound peptides were washed with 90 µL of ammonia water (pH 10) followed by elution with a series of ammonia water containing increasing concentration of ACN (6%, 9%, 12%, 15%, 18%, 21%, 25%, 30%, 35%, and 50%). The fractions with 25%, 30%, 35% and 50% of ACN were combined with fractions containing 6%, 9%, 12% and 21% of ACN, respectively. The resultant 6 fractions were the dried and stored at -80 oC for LC-MSn analysis.

### LC-MS^3^ analysis of DSSO cross-linked peptides

The SEC-HpHt fractions were subjected to LC MS^3^ analysis using an UltiMate 3000 RSLC nano-HPLC system coupled to an Orbitrap Fusion Lumos mass spectrometer (Thermo Fisher Scientific) as described previously^27^. The peptides were separated by RPLC (50 cm x 75 μm Acclaim PepMap C18 column, Thermo Fisher Scientific) with over an 87-min gradient of ACN (4% to 25%) at 300 nL/min flow rate. MS1 scans were measured in the Orbitrap with a scan range from 375 to 1800 m/z, 60,000 resolution, and AGC target 4×105 at top speed per 4 s cycle time. Ions with charge 4+ or greater were selected for MS2 and subjected to fragmentation using CID with NCE 23. For MS2 scans, the resolution was set to 30,000, AGC target 5e4, precursor isolation width 1.6 m/z, and maximum injection time 100 ms. A targeted inclusion on ions with mass difference corresponding to the difference in alkene and thiol DSSO fragments (31.9721 Da) was used to select precursors for MS3 analysis. For MS3 scans, HCD was used with a normalized collision energy of 28%, the AGC target was set to 2×104, and the maximum injection time was set to 125 ms.

### Identification of DSSO cross-linked peptides

Peaklists were extracted from the LC MSn raw files using the in-house software PAVA (UCSF) and the extracted MS3 spectra were searched against a SwissProt database (2021.10.02 version) concatenated with its randomized decoy sequences using Protein Prospector (v.6.3.5). The mass tolerances allowed were ±20 ppm for precursor ions and 0.6 Da for fragment ions. The database search was performed with trypsin as a protease with a maximum of three allowed missed cleavages. Cysteine carbamidomethylation was set as the fixed modification. The variable modifications included N-terminal protein acetylation, methionine oxidation, and N-terminal conversion of glutamine to pyroglutamic acid. Additionally, three specific modifications resulting from DSSO were included in the search: thiol (C3H2SO, +86 Da), alkene (C3H2O, +54 Da), and sulfenic acid (C3H4O2S, +104 Da)^23^. The in-house software XL-Tools was used to automatically identify, summarize and validate cross-linked peptides based on Protein Prospector database search results and MS^n^ data. No decoy hits were found after the integration of MS^1^, MS^2^ and MS^3^ data.

### Development of MS^3^ based XL-MS analysis tool

We developed Ving, a software to assess the MS^2^/MS^3^-based cleavable cross-linking database search results to produce a set of cross-linked-spectrum matches (CSMs) (**Extended Data Fig. 1a**). Ving input consists of raw spectral data in mzML format^54^, and database search results of MS^2^ and MS^3^ spectra in PepXML format^55^. The output of Ving is a human-readable text file listing the CSMs observed from the spectral data and database search results. Ving functions by first parsing the mzML spectral data file to create spectral groups (SGs) consisting of MS^2^ and MS^3^ events that are associated with a single precursor ion selection. Each SG specifies the scan numbers and retention times of the spectra contained within the group, as well as the precursor ion mass and charge states of the MS^2^ and MS^3^ events. Next, database search results from two separate searches of either the MS^2^ or MS^3^ scan events are added to each SG. The searches were performed using the TPP^26^ as described previously, and are used to assign peptide sequences and associated probabilities to the scan events. As the MS^2^ and MS^3^ database searches are performed independently, peptide sequences and probabilities are first assigned to the MS^2^ events in each SG, then peptide sequences and probabilities are assigned to the MS^3^ events in each SG. Both MS^2^ and MS^3^ database search results are required, as the MS^2^-based results are essential for assessing whether or not a SG is derived from a single peptide precursor ion, i.e. produces a single peptide-spectrum match (PSM), while the MS^3^-based results are essential to identify both peptides if a SG is interpreted to be a CSM.

Assessment of each SG to determine probable CSMs occurs after all peptide sequence assignments have been made to all MS^2^ and MS^3^ spectra within all the groups. A series of thresholds categorize each group into either PSMs, or various types of CSMs. First, the probabilities of the peptide sequence assignments of the MS^2^ scan events are evaluated, and all assignments with a probability > 0.8 are assigned the status of a single, non-linked PSM. If the sequence assignment also contains evidence for a modification mass of the hydrolyzed cross-linker on an internal lysine, it is further classified as a dead-end or mono-linked PSM. For SGs with MS^2^ assignments of probability below 0.8, the MS^3^ peptide assignments and probabilities are evaluated. If multiple MS^3^-level peptide sequence identifications were made with a probability > 0.8 and containing a lysine residue with a modification mass approximating the cross-linker cleavage product, those sequences are further evaluated as candidate CSMs. If the masses of the two peptide sequences plus the crosslinker summed together to match the mass of the original precursor ion, then the group is classified as a CSM. If none of the peptide sequences sum to the precursor mass, despite evidence of a modified lysine, then the SG is classified as Incomplete CSM. If the SG has only zero or one MS^3^-level peptide sequence with a probability > 0.8, the group is classified simply as Unknown PSM. Following classification of all SG, a simple summary report is presented to the user and the entirety of the results are exported to a human-readable, tab-delimited text file.

### LC-MS analysis of PhoX cross linked peptides

The PhoX cross-linked peptides samples were analyzed on a timsTOF Pro mass spectrometer (Bruker Daltronics) as described preiviously^56^. Briefly, peptides from each SEC fraction 9 – 24 (**Extended Data Fig. 1c**) were loaded on to the column operated using UltiMate 3000 RSLC nano-HPLC system (Thermo Fisher Scientific) and eluted peptides were analyzed with the timsTOF Pro mass spectrometer using CaptiveSpray source (Bruker Daltonics). Peptides were first trapped on a C18 precolumn (Acclaim PepMap 100, 300 μm × 5 mm, 5 μm, 100 Å) (Thermo Fisher Scientific) and eluted peptides were subsequently separated on a μPAC 50 column (PharmaFluidics) over 180 min with ACN gradient ramping up from 3% to 35%. During elution, the flow rate of the gradient changed from 900 to 600 nL/min for the first 15 min, followed by a constant flow rate of 600 nL/min. The column was then washed for 15 minutes with higher ACN concentration (35% to 85%, 85%, 85% to 3%, for 5 minutes each) at a flow rate of 600 nL/min.

For MS analysis with the timsTOF Pro mass spectrometer, the mobility-dependent collision energy ramping settings were 95 eV at an inversed reduced mobility (1/ k_0_) of 1.6 V s/cm^2^ and 23 eV at 0.73 V s/cm2. The collision energies were interpolated linearly between the two 1/k_0_ values and were kept constant above or below. TIMS scans were not merged and the target intensity per individual parallel accumulation serial fragmentation (PASEF) precursor ion was kept at 20,000. The range of each scan was kept between 0.6 and 1.6 V s/ cm^2^ with a ramp time of 166 ms. The number of PASEF MS/MS scans triggered were 14 per cycle (2.57 s) with a maximum of seven allowed precursors per mobilogram. The precursor ions selected for fragmentation ranged between *m/z* 100 and 1700 with charge states between 3+ to 8+. The active exclusion was allowed/set to 0.4 min (mass width 0.015 Th, 1/k_0_ width 0.015 V s/cm^2^).

### TimsTOF MS data analysis

TimsTOF-MS data were converted to .mgf format using MSConvert^57^. The mgf files were then processed for identification of cross-linked peptides using pLink-2(ref.^58^) with default settings unless stated otherwise. All files were searched against Nomo-1 surfaceome specific custom database generated from regular DDA analysis. The custom database was generated from SEC fractionated samples. For pLink based cross linked peptide analysis, trypsin was set as the protease allowing three missed cleavages. Cysteine carbamidomethylation was set as fixed modification with methionine oxidation and N-terminal acetylation as variable modification. The search was performed with ±20 ppm mass tolerance window for precursor as well as fragment ions, and results were reported at 1% FDR.

### Flow cytometry

Immunostaining of cells were performed as per the instructions from antibody vendor unless stated otherwise. Briefly, 1 million cells were resuspended in 100 µl of FACS buffer (PBS + 2% FBS) with 1 ug antibody added to it. The cells were incubated at 4C for 10-15 minutes and then washed thrice with the FACS buffer. For staining active form of ITGB2, antibody incubation step was performed at 37C for 1 hour. In case of staining primary AML cells for activated ITGB2, recipe of FACS buffer was RPMI-1640 + 5% FBS + 2% BSA + 50 µg/ml DNase-I (Gold Biotechnology, D-301-500). For all other primary cell staining, FACS buffer recipe was D-PBS + 5% FBS + 2% BSA + 5 mM EDTA + 50 µg/ml DNase-I with Human Trustain (Biolegend, 422302). All the compensation was done using UltraComp eBeads™ Compensation Beads (Invitrogen, 01-2222-42). All the flow cytometry analysis was done with Cytoflex (Beckman Coulter) and data was analyzed using FlowJo_v10.8.1. The antibodies used in this study are CD3 (Biolegend, 980008, 300412, clone-UCHT1), CD19 (Biolegend, 363006, 363036, clone-SJ25C1), CD45 (Biolegend, 368512, clone-2D1), CD14 (Biolegend, 367118, 367104, clone-63D3), CD34 (Biolegend, 343510, 343510, clone-581), CD69 (Biolegend, 985206, clone-FN50), CD11a/CD18 (Biolegend, 363406, 363416, clone-m24), CD18 (Biolegend, 302106, clone-TS1/18), CD33 (Biolegend, 303404, clone-WM53), CD62L (BD Biosciences, 559772, clone: DREG-56), CD45RA (Thermo Fisher Scientific, 12-0458-42, clone: HI100), CD16 (Biolegend, 302032, clone-3G8) and CD64 (Biolegend, 305018, clone-10.1). Secondary antibody used was anti-human IgG Fc antibody (Biolegend, 410720). All the respective isotype antibodies used were procured and used as per the vendor’s instructions.

### Phage display selections

A synthetic, phage-displayed Fab library^45^ was selected for binding to either Integrin-β2/Integrin αM (R and D 4047-AM, Antibody #7062, 7#063, #7065) or Integrin-β2/Integrin αL (R and D 3868-AV, Antibody # 7060, #7341) recombinant protein complexes. Briefly, Integrin-β2 recombinant protein complexes were immobilized on Maxisorp Immuno plates (ThermoFisher, 12-565-135) and used for positive binding selections with library phage pools that were first exposed to neutravidin coated wells to deplete nonspecific binders. After four rounds of binding selections, clonal phage was prepared and evaluated by phage ELISA and sequencing as described^45^.

### Antibody production

Antibodies were produced using the human Expi293 expression system (Thermo Fisher). Expi293 cells (in 2 mL volume) were transiently transfected with construct DNA using FectoPro transfection reagent (Polyplus Transfection, 101000014). Following 5-day expression period, antibodies were purified using rProteinA Sepharasoe (GE Healthcare) and stored in phosphate buffer (50 mM NaH_2_PO_4_, 75 mM Na_2_HPO_4_, 100 mM H_3_PO_4_, 154 mM NaCl).

### Bio-Layer Interferometry (BLI) binding assays

The binding of human Integrin-β2 antibodies was tested against three different Integrin-β2 complexes including Integrin-β2/Integrin αM (R and D 4047-AM), Integrin-β2/Integrin αX (R and D 5755-AX), and Integrin-β2/Integrin αL (R and D 3868-AV). To determine the binding kinetic parameters of the antibodies, BLI experiments were performed on an Octet HTX instrument (Sartorius) at 1000 rpm and 25°C. All proteins were diluted in an assay buffer (PBS, 1% BSA, 0.05% Tween 20). Tested and negative control antibodies at a concentration of 2 µg/ml were first captured on AHQ biosensors to achieve the binding signals of 0.8-1.3 nm. Unoccupied Fc-binding sites on the antibody-coated sensors were subsequently quenched by 20 µg/mL of the Fc protein. After equilibration with the assay buffer, the biosensors were then dipped for 600 s into wells containing 5-fold serial dilution of Integrin-β2 complexes (association phase), followed by a transfer back into an assay buffer for additional 600 s (dissociation phase). Assay buffer alone served as a negative control. Binding response data were reference subtracted and were globally fitted with 1:1 binding model using ForteBio’s Octet Systems software v9.0.

### Non-specific ELISA panel

The ELISA protocol to assess interactions of the antibodies with unrelated macromolecules were performed as described previously^59^. The tested antigens included Cardiolipin (50 μg/mL, Sigma C0563), KLH (5 μg/mL, Sigma H8283), LPS (10 μg/mL, InvivoGen tlrl-eblps), ssDNA (1 μg/mL, Sigma D8899), dsDNA (1 μg/mL, Sigma D4522), and Insulin (5 μg/mL, Sigma I9278). In addition, the binding of each antibody was also tested against empty wells (BSA only control) and wells containing goat anti-human Fc antibody (positive control, 1 µg/mL, Jackson 109-005-098). The antigens were coated at 30 µL per well in 384-well Maxisorp plates and incubated at 4°C overnight. Plates were blocked with 0.5% bovine serum albumin (BSA) for 1 hour at room temperature and washed with PBS + 0.05% Tween20. The antibodies were added at 100 nM and allowed to bind for 60 min at room temperature. Plates were washed with PBS + 0.05% Tween20 and binding was detected with anti-kappa HRP antibody (1:5000, Southern Biotech #2060-05) and developed with the TMB substrate (KPL (Mandel) KP-50-76-03).

### Plasmid constructs

All the plasmid constructs were generated using NEBuilder® HiFi DNA Assembly Master Mix (NEB, E2621L) as per the vendor’s instructions with some modifications. The DNA fragments containing the binder (scFv) sequence along with the 40 bp vector compatible flanking region for Gibson assembly was procured from Twist Bioscience. Meanwhile, the target CAR plasmid backbone was linearized with BamHI-HF (NEB, R3136T) and cleaned up using Zymo Research DNA purification kit (Zymo Research, D4013). 10 ng of linearized vector and 5 ng of the DNA fragment (insert) was used to set 10 uL of gibson assembly reaction. This reaction mixture was then transformed into stbl3 competent *E.coli* cells (QB3 MacroLab, UC Berkeley) and the colonies obtained were screened for the positive clones with sanger’s sequencing services from Genewiz.

### Primary T cell isolation

Primary T cells were isolated from LeukoPaks obtained from Stem Cell Technologies (200-0092). CD8 and CD4 cells were isolated separately using their EasySep™ Human CD8+/CD4+ T Cell Isolation Kit as per manufacturer’s instructions. Briefly, all the unwanted cells were labelled with magnet conjugated antibody cocktail which is separated using their EasySep magnetic stand leaving CD4 or CD8 cell in suspension using the vendor supplied EasySep Human CD4/CD8 T Cell Iso Kit (Stem Cell Technologies, 17952 for CD4 and 17953 for CD8). This negative selection approach results in isolation of untouched CD8 or CD4 T cells and stored frozen with 10% DMSO (MP Biomedicals, 196055). In total, primary T cells from five different donors were used for the *in vitro* and *in vivo* studies here.

### CAR-T generation

T cells were thawed and grown in T cell media constituting Optmizer CTS media (Gibco, A10221-01) + CTS supplement (Gibco, A10484-02) + 5% Human AB Serum (Valley Biomedical, HP1022) + Penicillin/Streptamycin + glutamax (Gibco, 35050-061). Recombinant human IL7 (Peprotech, 200-07) and IL15 (Peprotech, 200-15), 10 ng/mL final concentration for each was freshly added to the cells every 2-3 days. For manufacturing CAR-T cells, primary T cells (CD4 or CD8) were thawed and cultured overnight. For the aITGB2 CAR-T, the cells were then additionally nucleofected with ribonuclease complex of ITGB2 sgRNA and Cas9 using P3 Primary Cell 4D-Nucleofector™ X Kit S (Lonza, V4XP-3032) using 4D-Nucleofector (Lonza) with its inbuilt program EO-115. The cells were then stimulated with 20 µl of CD3/CD28 Dynabeads (Thermo Fisher Scientific, 11131-D) per million cells. Meanwhile, lentivirus carrying the CAR expression cassette was added to the cells the day after adding the stimulation beads. The virus was withdrawn from the culture after 24 followed by 2-3 rounds of PBS wash using centrifugation at 300 RCF for 5 minutes. Assuming bead stimulation as day 0, beads were withdrawn on day 4 using magnetic rack and the T cells. On day 6 or 7, the cells were MACS sorted for the CAR positive cells using myc tag of the CAR constructs as a handle using biotinylated c-myc antibody (Milteni Biotec, 130-124-877). The CAR-T cells were used for *in vitro* and *in vivo* studies within day 10 – 14 of the manufacturing process.

### T cell activation assay

PBMC cells were treated with 3 µM ionomycin (Sigma Aldrich, 407950) + 25ng/mL LPS (Sigma Aldrich, L4391) + 100U/mL IL-2 (Prospec, CYT-209) and cultured overnight in CO2 incubator. The cells were then co-stained with CD3 and CD69 and analyzed with flowcytometry. CD3 was used to gate on T cells and CD69 was used as a T cell activation marker.

### In vitro cytotoxicity assay

The AML cell lines used for *in vitro* cytotoxicity analysis were engineered to stably express luciferase using lentiviral transduction. The cell lines were co-cultured overnight with CAR-T cells in various ratios in a 96 well white plate. 150 μg/mL of d-luciferin (Gold Biotechnology, LUCK-1G) was then added to each well and incubated for 3 - 5 minutes at RT, after which the plate is read for luciferase signal using GloMax Explorer Plate Reader (Promega). For each ratio (CAR-T : Tumor), the bioluminescence reading from the tumor cells co-cultured with untransduced T cells were considered 100% viable and thus used for normalization.

### Degranulation assay

CAR-T cells were co-cultured with tumor at ratio of 2:1 for 6 hours at 37o C in CO2 incubator with CD107a antibody (Biolegend, 328620, clone-H4A3) and golgistop (BD Biosciences, 51-2092KZ). The cells were washed twice with centrifugation at 500 RCF for minutes at RT. Levels of CD107a was then measured with flow cytometer as a read out of degranulation. CAR-T cells were labelled with GFP which was used for gating them for analysis.

### Generation of ITGB2 knockout cells

Knockout cell lines or Primary T cells were generated using invitro nucleofection of Cas9 ribonuclease protein complex. Briefly, 2 μl each of sgRNA (100 µM) (Synthego Corporation) and recombinant Cas9 protein (40 µM) (QB3 MacroLab, UC Berkeley) was incubated at 37° C for 15 minutes. The sgRNA used in this study were obtained from Brunello library^60^ (sgRNA-1-TCAGATAGTACAGGTCGATG, sgRNA-2-CTCCAACCAGTTTCAGACCG, sgRNA-3-TCAGGGTGCGTGTTCACGAA, sgRNA-4-TCATCCCCAAGTCAGCCGTG). Meanwhile, 1e6 cells were washed once with PBS (500 x *g* for 5 min at RT) and resuspended in a mixture of 16.4 µl SF cell line solution and 3.6 µl supplemental solution-1 (Lonza, V4XC-2032). The sgRNA and Cas9 ribonuclease protein complex under incubation was then mixed with the cell suspension and 20 µl of it was put a cuvette and nucleofected using 4D-Nucleofector (Lonza) with the inbuilt program DS-137 for cell lines and EO-115 for primary T cells. The cells were then allowed to rest at RT for 2-5 minutes and added to fresh media pre-warmed at 37° C.

### Humanized Immune System (HIS) mice generation

All the mice used for HIS mice generation were of NSG-SGM3 strain (NOD.Cg-*Prkdc^scid^ Il2rg^tm1Wjl^* Tg(CMV-IL3,CSF2,KITLG)1Eav/MloySzJ) and obtained from Jackson Laboratories. Each mouse was treated with busulfan (12.5 mg/kg) for two consecutive days then one recovery day followed by injection with 70,000 CD34+ human hematopoietic cells intravenously through tail vein. Fully de-identified human CD34 cells enriched blood samples were obtained from Bone Marrow and Transplantation Laboratory at UCSF and were MACS sorted using CD34 MicroBead Kit (Miltenyi Biotec, 130-046-702) and incubated with CD3 antibody (Biolegend, 317302, clone – OKT3) for T cell depletion10 minutes prior to injection in mice, to limit any possible development of graft-vs-host disease. The blood draw of these mice were analyzed using flow cytometry 8 weeks post CD34 cells injection, to determine the engraftment efficiency using Human CD45+ cells as a read out (>1.5% threshold).

### Murine CAR-T efficacy experiments

All the mice used in the experiments were 6-8 weeks old (either all male or all female for a particular study) and obtained from either Jackson laboratory (NSG-SMG3) or from in-house (NSG) bred stocks of Pre-Clinical Therapeutics Core of UCSF. Each mouse was injected with 1 million AML cell lines or 2 million PDX AML lines intravenously through tail vein. In case of PDX, the mice were irradiated with dose of 250cGy 4-6 hours prior to injection. 5 days later, each mice were treated with a total of 5 million CAR-T cells at 1:1 ratio of CD4 and CD8.

Tumor burden in case of cell lines (luciferased lines) was assessed using bioluminescence imaging with Xenogen In Vivo Imaging System (Caliper Life Sciences). In case of PDX, using flowcytometry analysis of blood draws and spleen size determination with ultrasonography was used as a readout for tumor burden. All the mice experiments were conducted in accordance with UCSF Institutional Animal Care and Usage Committee.

### Statistical analysis

All statistical analysis was performed using GraphPad Prism v.9 unless stated otherwise. The data have been represented as ± mean and *p*-value < 0.05 were considered statistically significant. All the proteomics related statistics were performed by the respective analysis suite used and stated in those sections. All mice were randomized before therapeutics treatment. Other statistical details are stated in the legends of the respective figures.

### Data Availability

Raw proteomic data generated here has been deposited at the ProteomeXchange/PRIDE repository with accession numbers: PXD035404, PXD035589 and PXD035591.

[Reviewer access details: Username: Password: xXqWy1wG]

[Reviewer access details: Username: Password: KYp5TVLG]

[Reviewer access details: Username: Password: foasTSFF]

### Code Availability

Ving software package for analysis of DSSO XL-MS data is available at github - https://github.com/mhoopmann/Ving.

---

**Supplementary Table 1.** Binding affinities (KD) of phage-display generated antibody clones against recombinant ITGB2 heterodimers. Obtained from biolayer interferometry; values based on curve fits as in Octet Systems software, as in Fig. 3c.

| Antibody ID | BLI K<sub>D</sub> (M) |  |  |
| --- | --- | --- | --- |
|  | ITGAL-B2 | ITGAM-B2 | ITGAX-B2 |
| 7060 | 2.16E-09 | 2.01E-09 | 1.51E-09 |
| 7062 | 2.91E-09 | 3.20E-09 | 1.46E-09 |
| 7063 | 1.02E-08 | 1.14E-08 | 3.96E-09 |
| 7065 | 1.56E-09 | 2.15E-09 | 2.03E-09 |
| 7341 | 4.07E-09 | 2.75E-09 | 2.80E-09 |

**Extended Data Figure 1.**
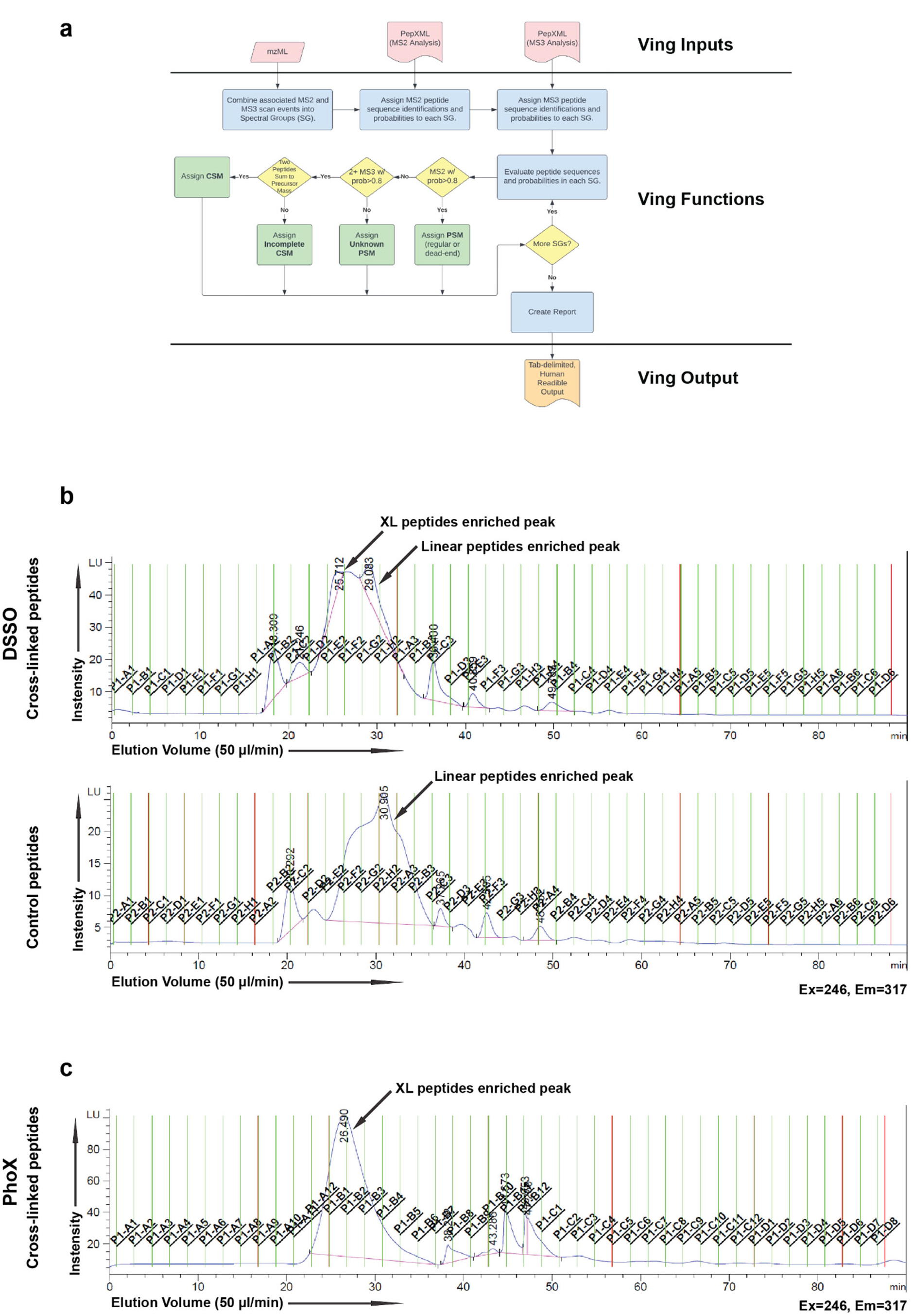
Ving and XL-MS SEC. **a.** Schematic workflow describing the working principle of Ving. **b.** Representative SEC trace of peptides obtained from DSSO cross-linked samples. **c.** Representative SEC trace of peptides obtained from PhoX cross-linked samples. For both strategy (DSSO and PhoX), samples were processed in 4 separate batches and every time SEC trace pattern were similar. XL peptides refers to cross-linked peptides.

**Extended Data Figure 2.**
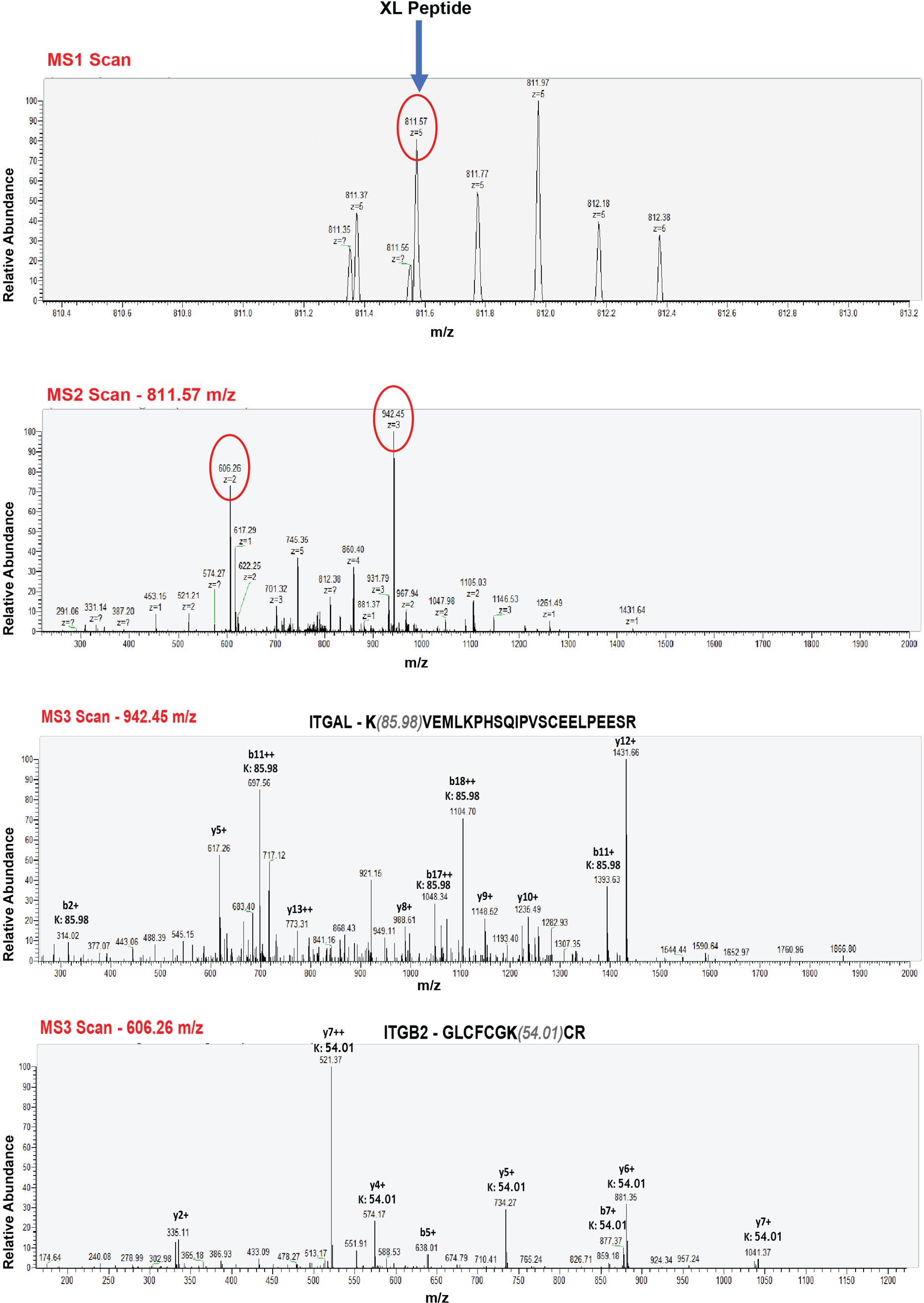
Representative MS spectra demonstrating the MS^3^ based strategy of XL-MS. Representative MS spectra demonstrating the MS^3^ based strategy of XL-MS. The Cross-linked peptides with 811.57 m/z gets selected for MS2 where the cross-linker gets cleaved in collision cell generating two separate peptides – 606.26 m/z and 942.45 m/z. These two high abundant peptides are then selected for MS^3^ where they undergoes full fragmentation for peptide identification; where we also note the respective modification on Lysine residues resulting from the cross-linker.

**Extended Data Figure 3.**
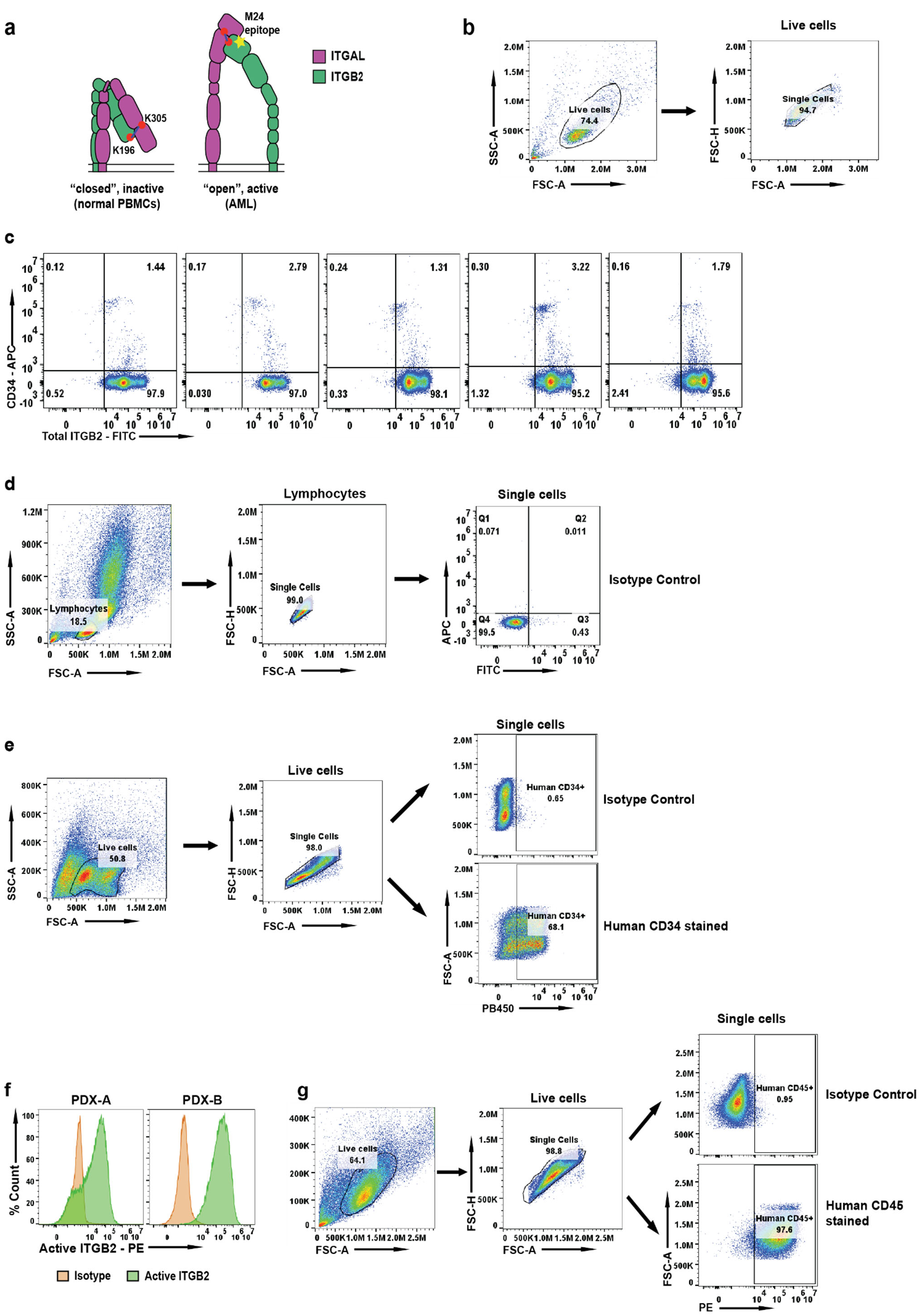
Discerning active integrin-β2 expression. **a.** Cartoon diagram showing proposed inactive and active conformations of ITGB2. **b.** Flow cytometry gating strategy for (Fig 2b, Fig 3d, Fig 4d). **c.** Flow cytometry plot showing presence of total Integrin-β2 on CD34+ HSPCs from GM-CSF mobilized peripheral blood. Cells were gated on singlet cells for analysis. Deidentified patient samples were used for this analysis (*n* = 5, independent donors). Representative of 1-2 independent experiments. **d.** Flow cytometry gating strategy for (c) and (Fig. 2c). **e.** Flow cytometry gating strategy for (Fig. 2d). **f.** Flow cytometry analysis showing expression of active ITGB2 in PDX models of AML (PDX-A and PDX-B). The y-axis represents percent count normalized to mode. Cells were gated on human CD45+ population cells for analysis. Representative plot from n = 2 separate PDX models of AML. **g.** Flow cytometry gating strategy for (f).

**Extended Data Figure 4.**
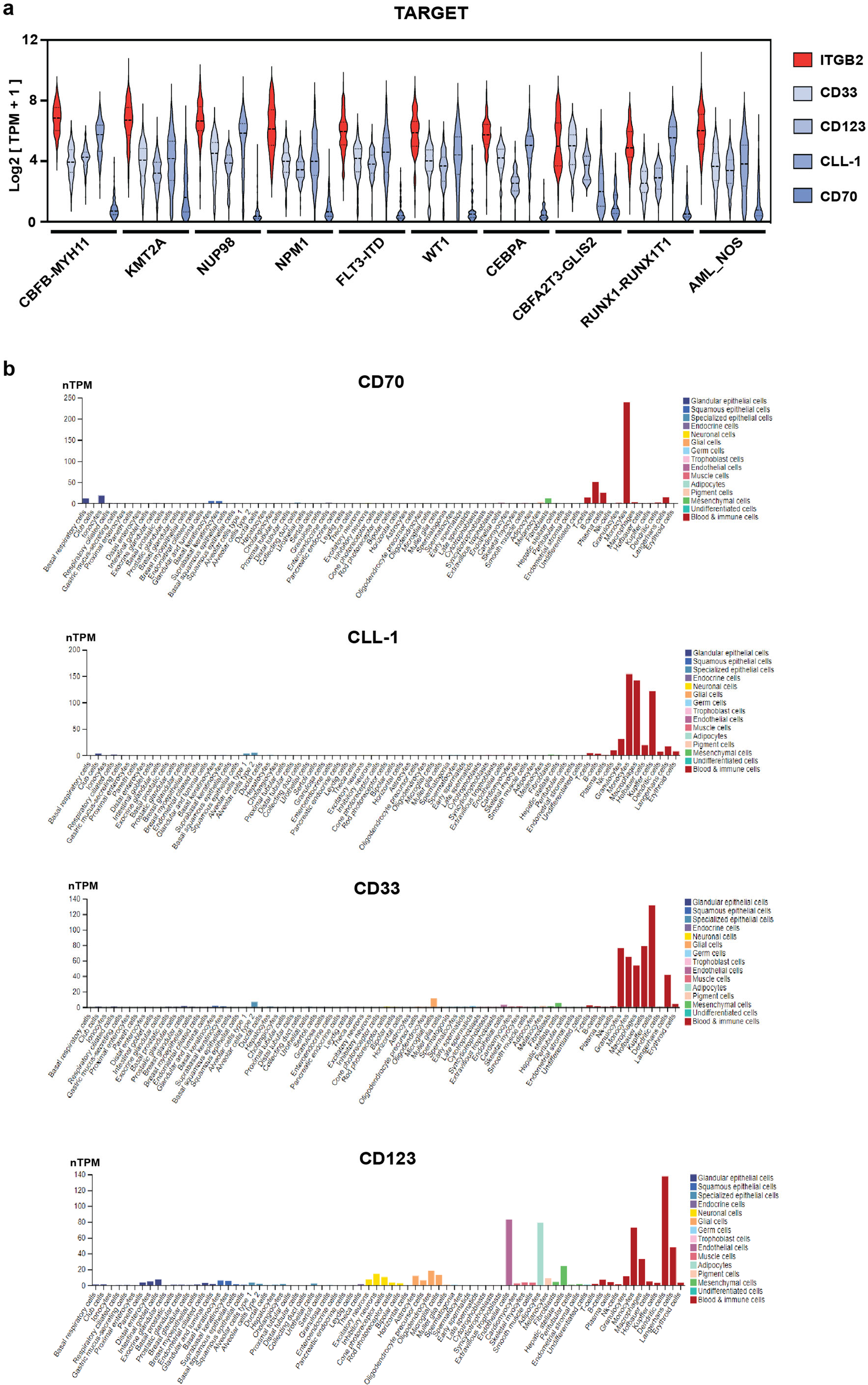
*ITGB2* transcript expression. **a.** AML subtype specific expression analysis of *ITGB2* and other notable AML targets of patient samples from TARGET database. **b.** Single cell sequencing data showing expression of notable AML target across normal human tissues and immune cells (adapted from Human Protein Atlas^42^).

**Extended Data Figure 5.**
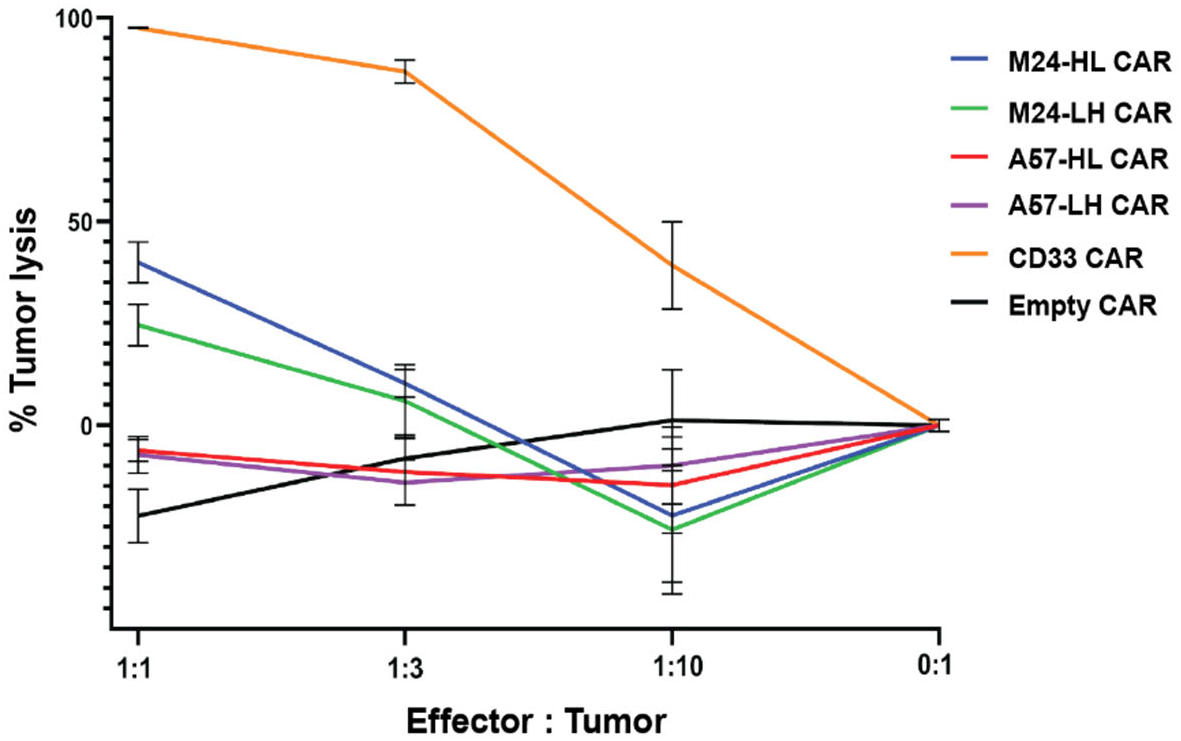
Initial anti-active integrin-β2 CAR-T designs. Luciferase based cytotoxicity analysis of M24 and A57 antibody derived CAR-T cells against AML cell line Nomo-1. The experiment was performed with 3 technical replicates in 2 independent experiments. The luciferase signals of the cytotoxicity assays were normalized against untransduced CAR-T of their respective E:T ratios. All the statistical data in this figure are represented as mean ±SEM.

**Extended Data Figure 6.**
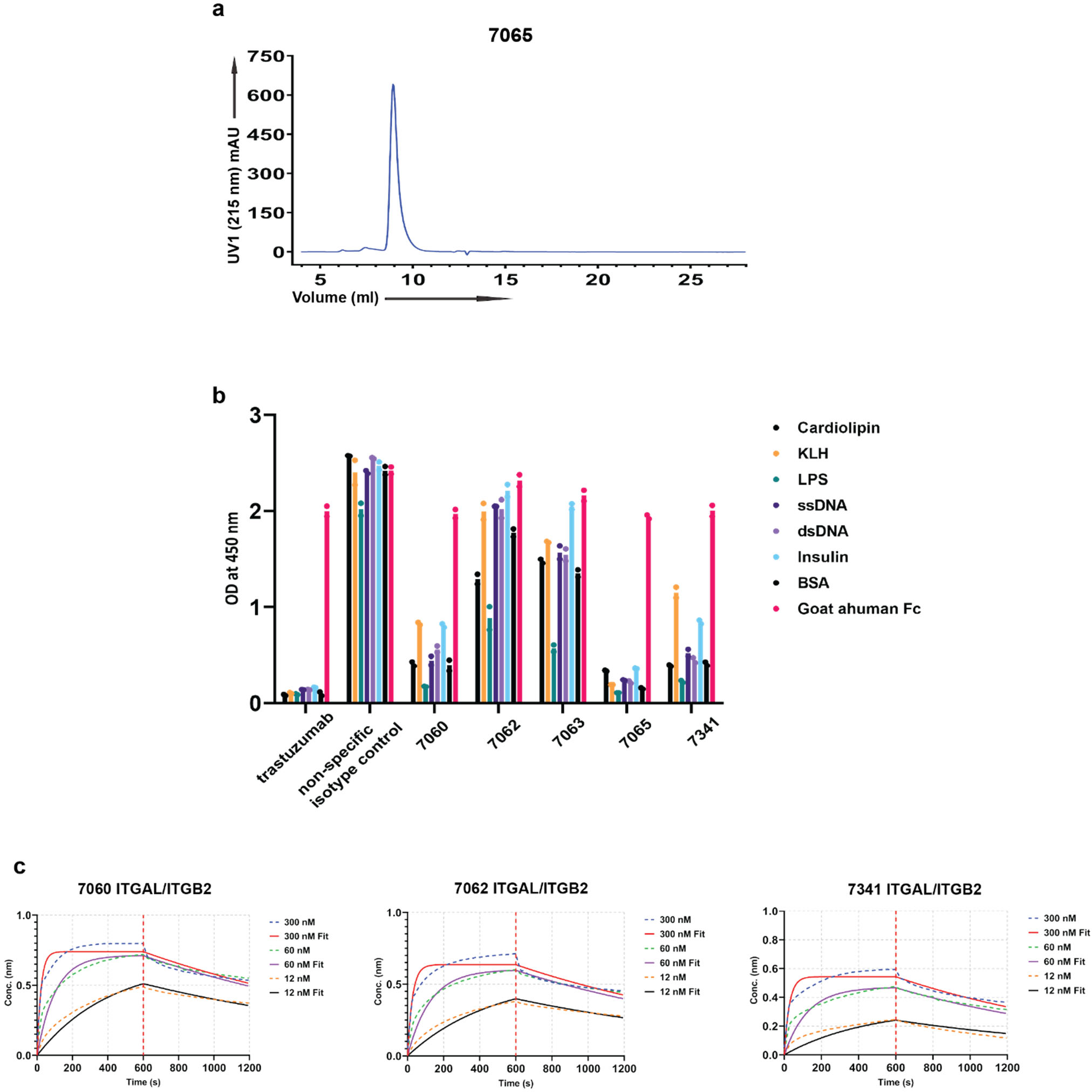
Characterizing recombinant antibodies to integrin-β2. **a.** Representative SEC trace of the antibody 7065 showing distinct peak, for quality check. **b.** Non-specific ELISA panel showing specificity profiles of the antibodies obtained from phage display selection. Experiment performed with *n* = 2 technical replicates. **c.** Representative BLI plots showing binding affinities (KD) of the antibodies against ITGB2 with their alpha partners. Each experiment was performed with *n* = 3 different concentrations of antibody.

**Extended Data Figure 7.**
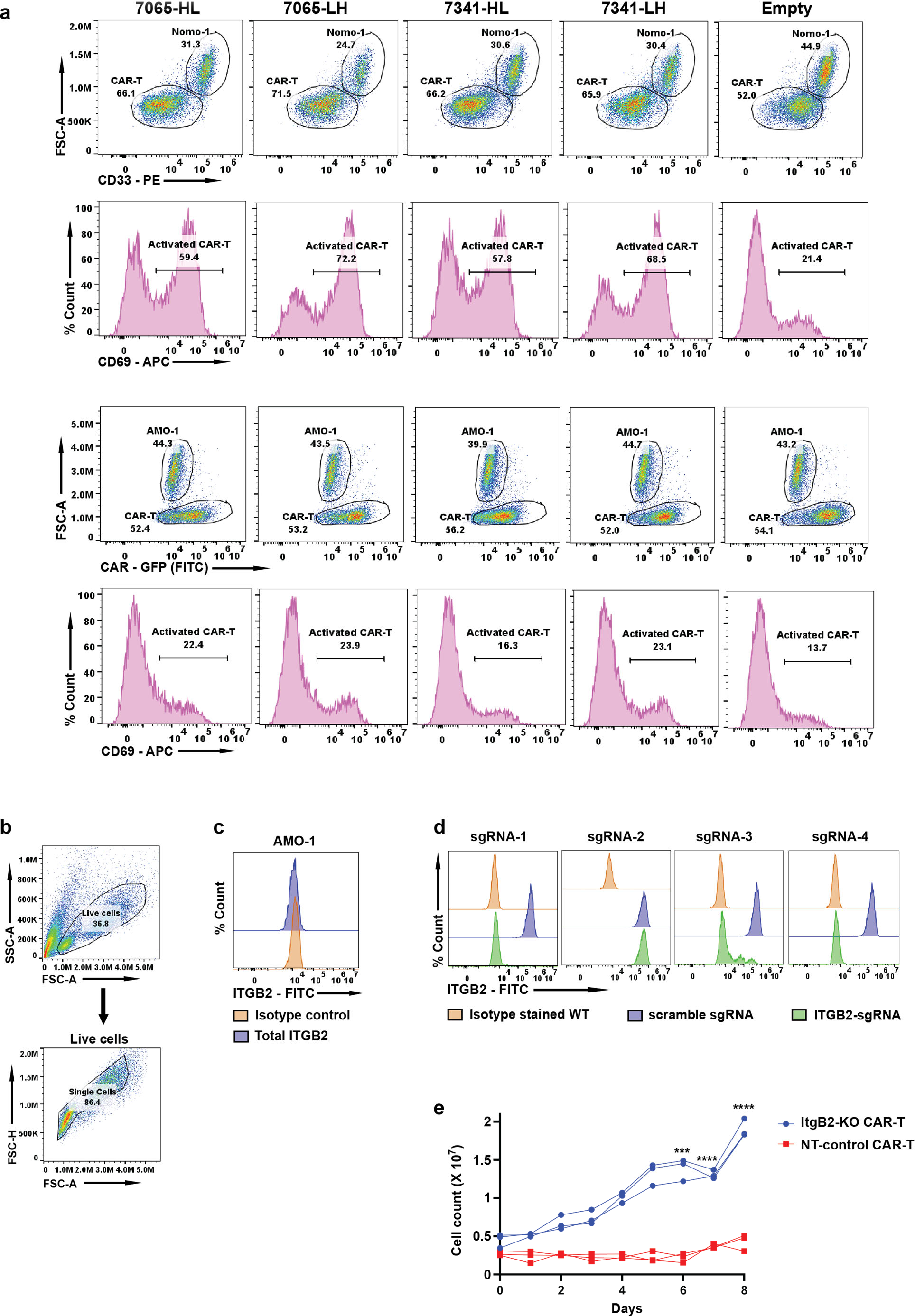
Evaluation of aITGB2 CAR-T designs incorporating recombinant antibodies. **a.** Flow cytometry screen for cytotoxicity and activation status of aITGB2 CAR-T designs vs. AML cell line Nomo1 (*n* = 1 for each design). Similarly, as a demonstration of specificity, cytotoxicity and activation status was also checked against AMO-1 (multiple myeloma cell line that does not express integrin-β2). Cells were gated on single cells for analysis. **b.** Flow cytometry gating strategy for (a). **c.** Flow cytometry analysis showing absence of ITGB2 in AMO-1 (*n* = 1). Cells were gated on single cells for analysis. Flow cytometry gating strategy similar to shown in (Extended Data Fig.3b). **d.** Flow cytometry analysis showing knockout efficiency of the various sgRNA used for knocking out ITGB2 in primary T cells. (*n* = 1 for each sgRNA). Cells were gated on single cells for analysis. Flow cytometry gating strategy similar to shown in (Extended Data Fig.3b). **e.** Plot showing proliferation of aITGB2-CAR-T cells, “with ITGB2 knockout” vs “Non-Targeting (NT) – control”. Scrambled sgRNA was used for NT-control. n=3 technical replicates. *p*<0.005 by two-tailed t-test at points noted.

**Extended Data Figure 8.**
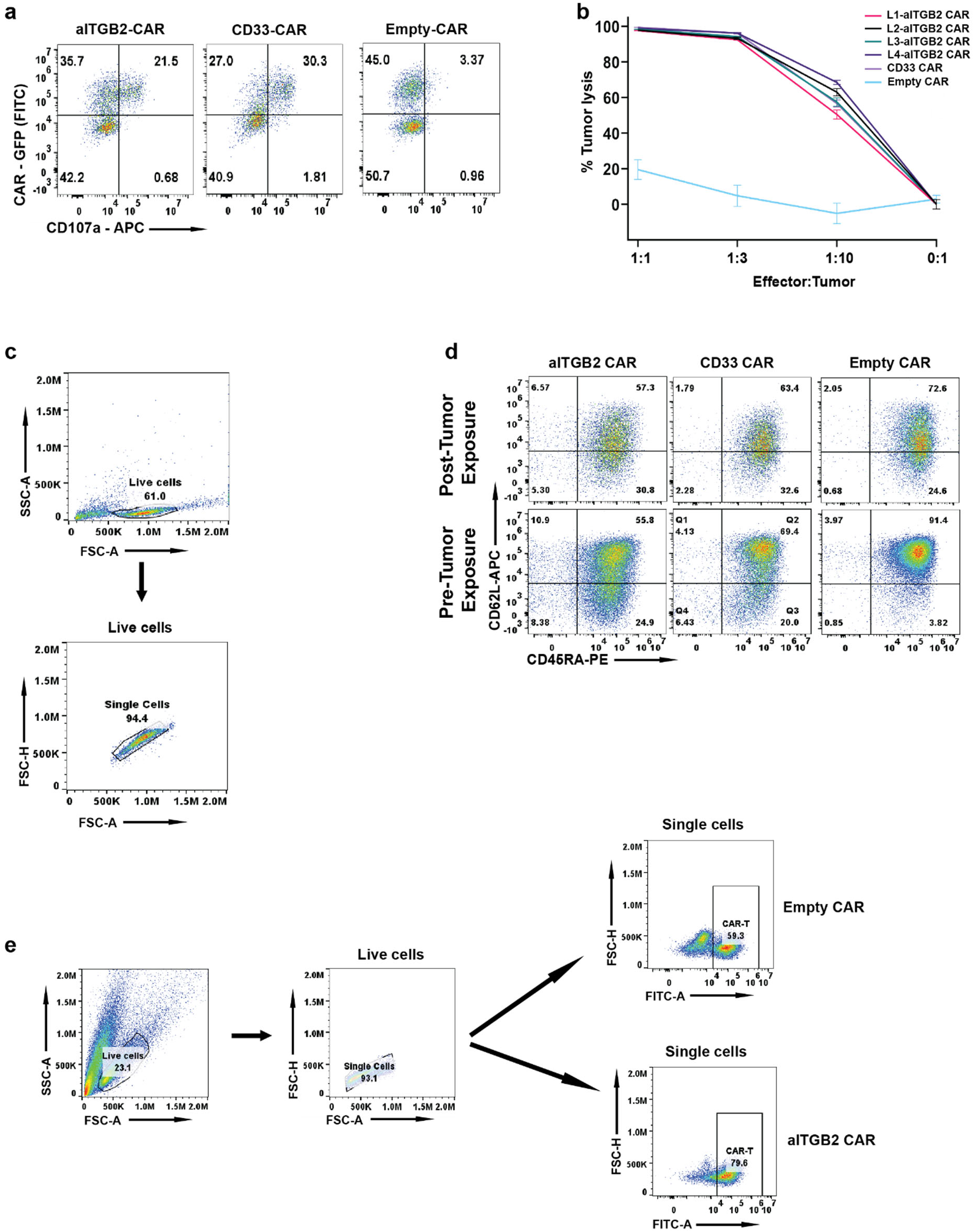
Additional aITGB2 evaluation. **a.** Degranulation assay of aITGB2 and anti-CD33 CAR-T against Nomo1 based on CD107a staining. CAR positivity denoted by GFP tag on CAR construct. E:T ratio was 1:1 and 6 hours incubation time (*n* = 1). Cells were gated on single cells for analysis. Flow cytometry gating strategy similar to shown in (Extended Data Fig. 7b). **b**. Luciferase assay-based cytotoxicity analysis showing efficacy of 7065 based aITGB2 CAR-T with 1X - 4X Gly4Ser (L1-L4) linker between heavy and light chain. *n* = 3 technical replicates; representative plot from 3 independent experiments. The luciferase signals of the cytotoxicity assays were normalized against untransduced CAR-T of their respective E:T ratios. **c**. Flow cytometry gating strategy for (Fig. 5d) **d.** Memory marker (CD45RA and CD62L) analysis of aITGB2 CAR-T and its comparison with CD33 CAR-T post tumor exposure at E:T ratio of 1:1 for overnight incubation. Upper right quadrant indicates naïve-like phenotype (*n* = 1). Cells were gated on single cells for analysis. **e.** Flow cytometry gating strategy for (d). All the statistical data in this figure are represented as mean ±SEM.

**Extended Data Figure 9.**
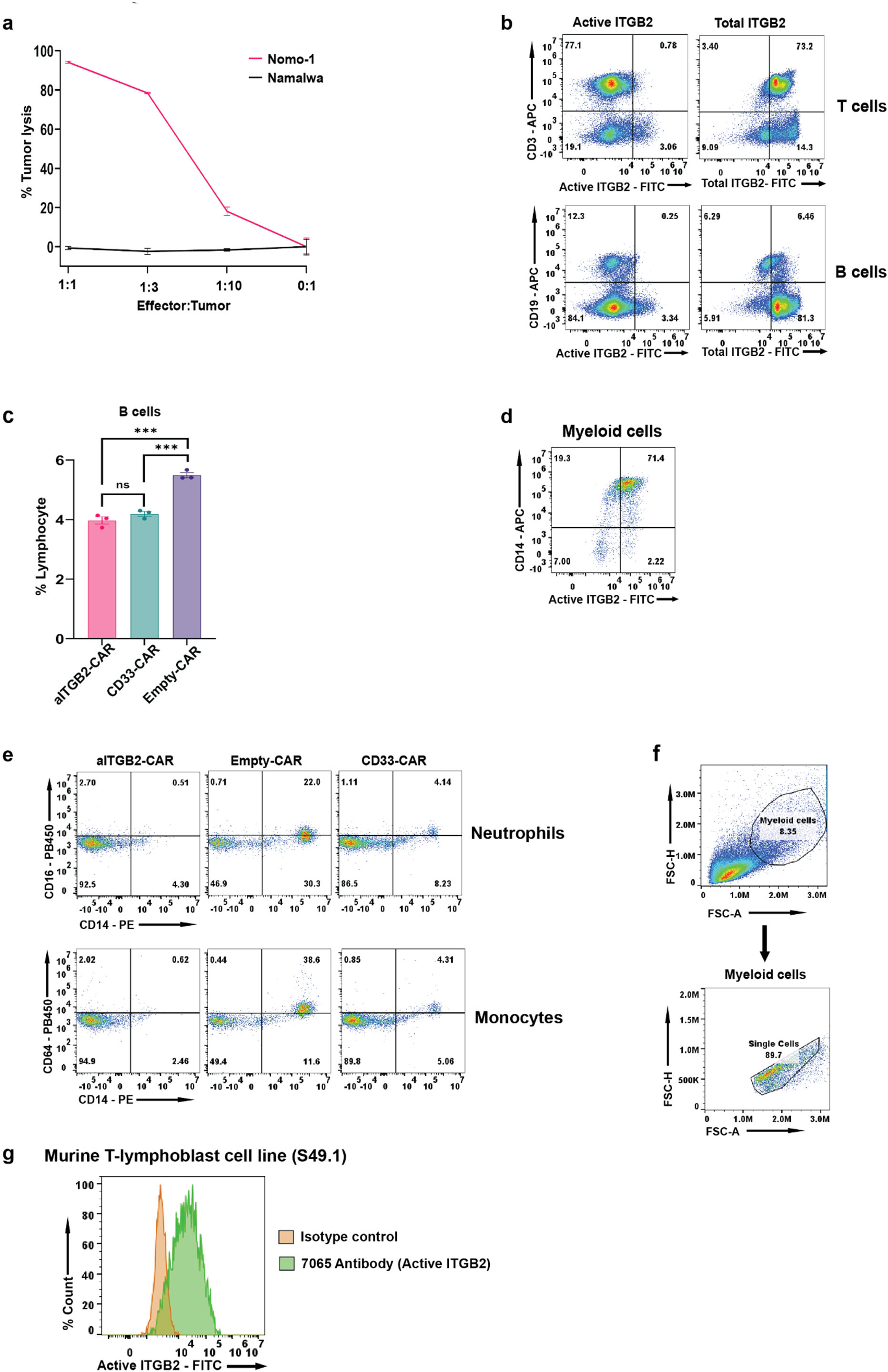
Determining specificity of aITGB2 CAR-T. **a.** Luciferase assay-based cytotoxicity analysis showing no activity of aITGB2 CAR-T vs. Namalwa (B-ALL) line) which does not harbor active ITGB2 although it does have total form of ITGB2 (see Fig. 2b). Nomo-1 as the positive control (*n* = 3 technical replicates). The luciferase signals of the cytotoxicity assays were normalized against untransduced CAR-T of their respective E:T ratios. All the statistical data in this figure are represented as mean ±SEM. **b.** Representative flow cytometry analysis showing absence of active ITGB2 and presence of total ITGB2 in T and B cells (*n* = 3 independent experiments). Cells were gated on single cells for analysis. Flow cytometry gating strategy similar to shown in (Extended Data Fig. 3d.) **c.** Flow cytometry analysis showing non-specific depletion of B cells with aITGB2 and anti-CD33 CAR-T (*n* = 3 technical replicates). Representative data from 3 independent experiments. *p*-value by two-tailed *t*-test. **d.** Representative flow cytometry analysis showing presence of active ITGB2 in myeloid cells (*n* = 3 independent experiments). Cells were gated on single cells for analysis. Flow cytometry gating strategy similar to shown in (f). **e.** Flow cytometry analysis showing cytotoxicity of aITGB2-CAR against neutrophils and monocytes in vitro (*n* = 2 PBMC donor). Cells were gated on single cells for analysis. **f.** Flow cytometry gating strategy for (d) and (e) **g.** Representative flow cytometry analysis showing cross reactivity of 7065 antibody against the murine ITGB2 on S49.1 cell line (*n* = 3 independent experiments). The y-axis represents percent count normalized to mode. Cells were gated on single cells for analysis. Flow cytometry gating strategy similar to shown in Extended Fig. 3b. All the statistical data in this figure are represented as mean ±SEM.

**Extended Data Figure 10.**
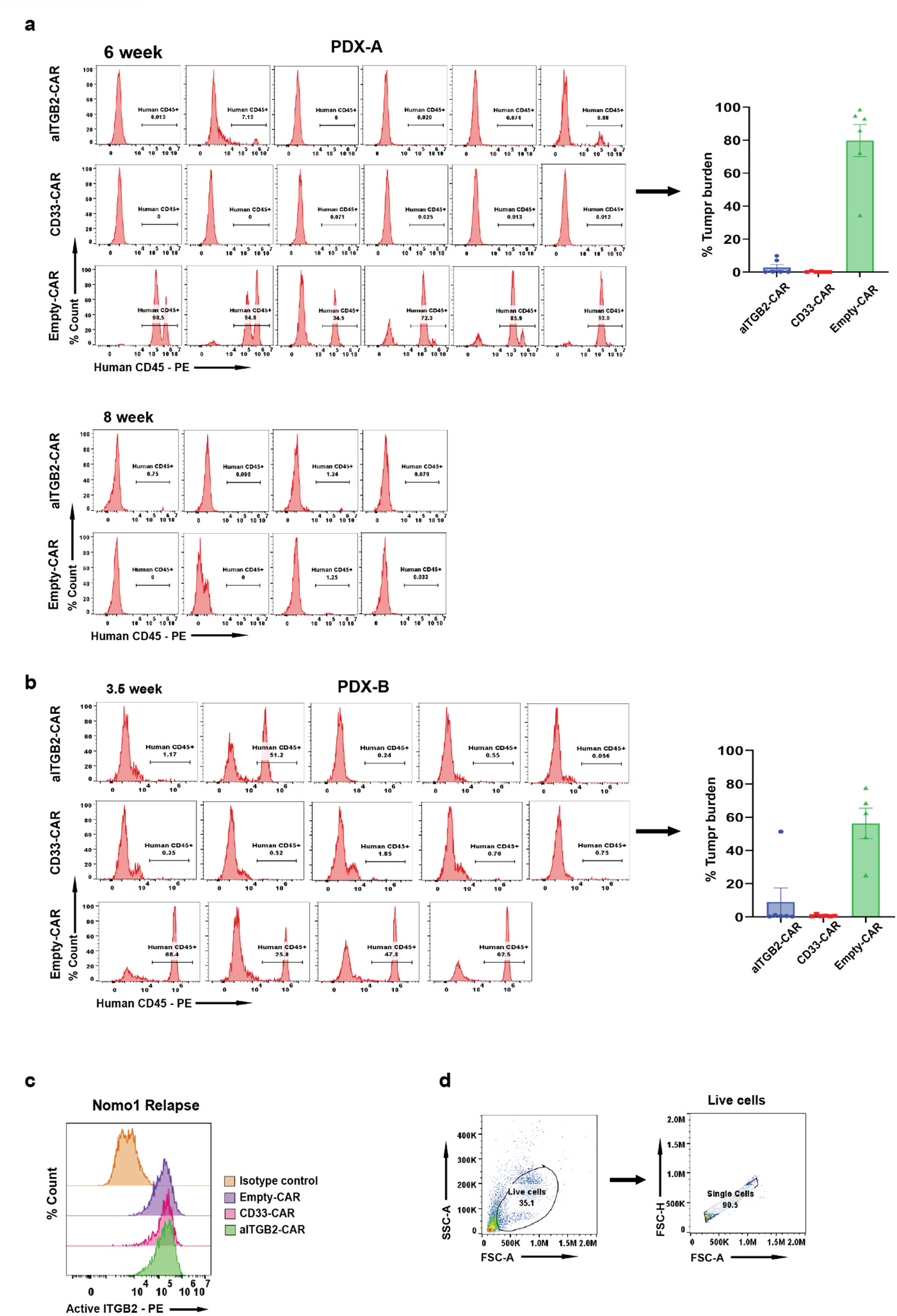
aITGB2 efficacy in PDX models. **a.** Flow cytometry analysis and bar graph of peripheral blood draw showing tumor burden at 6 and 8 weeks post tumor injection of PDX-A. The y-axis represents percent count normalized to mode. Cells were gated on single cells for analysis. Representative of data from *n* = 4 - 6 mice per arm dependent on number of mice was alive until designated time point. **b.** Flow cytometry analysis and bar graph of peripheral blood draw showing tumor burden at 3.5 weeks post tumor injection of PDX-B. The y-axis represents percent count normalized to mode. Cells were gated on single cells for analysis. Representative of data from *n* = 5 - 6 mice per arm dependent on number of mice was alive until designated time point. **c.** Flow cytometry analysis showing active ITGB2 density of tumor cells harvested from relapse Nomo-1 mice model (*n* = 1 mouse per condition). The y-axis represents percent count normalized to mode. Cells were gated on human CD45+ cells for analysis. Flow cytometry gating strategy similar to shown in (Extended Data Fig. 3g). **d.** Flow cytometry gating strategy for (a) and (b). All the statistical data in this figure are represented as mean ±SEM.

